# *ctQC* improves biological inferences from single cell and spatial transcriptomics data

**DOI:** 10.1101/2024.05.23.594978

**Authors:** Vairavan Lakshmanan, Merve Kahraman, Dominique Camat Macalinao, Nicole Gunn, Prasanna Nori Venkatesh, Chang Meihuan, Cherylin Fu, Leow Wei Qiang, Iain Beehuat Tan, Shyam Prabhakar

## Abstract

Quality control (QC) is the first critical step in single cell and spatial data analysis pipelines. QC is particularly important when analysing data from primary human samples, since genuine biological signals can be obscured by debris, perforated cells, cell doublets and ambient RNA released into the “soup” by cell lysis. Consequently, several QC methods for single cell data, employ fixed or data-driven quality thresholds. While these approaches efficiently remove empty droplets, they often retain low-quality cells. Here, we propose cell type-specific QC (*ctQC*), a stringent, data-driven QC approach that adapts to cell type differences and discards soup and debris. Evaluating single cell RNA-seq data from colorectal tumors, human spleen, and peripheral blood mononuclear cells, we demonstrate that *ctQC* outperforms existing methods by improving cell type separation in downstream clustering, suppressing cell stress signatures, revealing patient-specific cell states, eliminating artefactual clusters and reducing ambient RNA artifacts. When applied to sequencing-based spatial RNA profiling data (Slide-seq), *ctQC* improved spatial coherence of cell clusters and consistency with anatomical structures. These results demonstrate that strict, data-driven, cell-type-specific QC is applicable to diverse sample types and substantially improves the quality and reliability of biological inferences from single cell and spatial RNA profiles.

## Introduction

Single cell RNA-seq (scRNA-seq) is a well established and widely used technique for defining cell types and states, identifying disease biomarkers and mechanisms, characterizing developmental dynamics and response to perturbations, inferring gene regulatory mechanisms and cell signalling interactions, and interpreting results from genetic and environmental association studies [1–5]. However, scRNA-seq analysis is vulnerable to data quality artifacts, which can distort or obscure the underlying biology [6, 7]. For example, dissociating single cells from solid tissues can induce apoptosis or necrosis, trigger stress responses, compromise cell membrane integrity, cause transcript degradation, contaminate the buffer solution with leaked RNA (ambient RNA) and fragment cells (debris) [6, 8–10] (Fig.1A). Freeze-thaw cycles and long storage times also compromise scRNA-seq data quality. Incomplete dissociation or loading at high cell concentration can introduce doublets, i.e. transcriptomes that represent more than one cell [11]. These artifacts are exacerbated when sample quality is low. Sample quality is a particular concern for primary human tissues, due to ischemia during surgery and delays in sample transfer from clinic to bench [12]. Finally physical and biochemical changes during scRNA-seq library preparation protocols may diminish data quality. To minimize data quality artifacts and preserve the integrity of downstream biological conclusions, virtually all scRNA-seq data analysis pipelines include an initial quality control (QC) step that removes low-quality cells [13, 14]. In addition to cell QC, multiple algorithms exist for characterizing ambient RNA and correcting the gene expression matrix for example SoupX [10], decontx [15] and cellbender [16].

**Figure 1:**
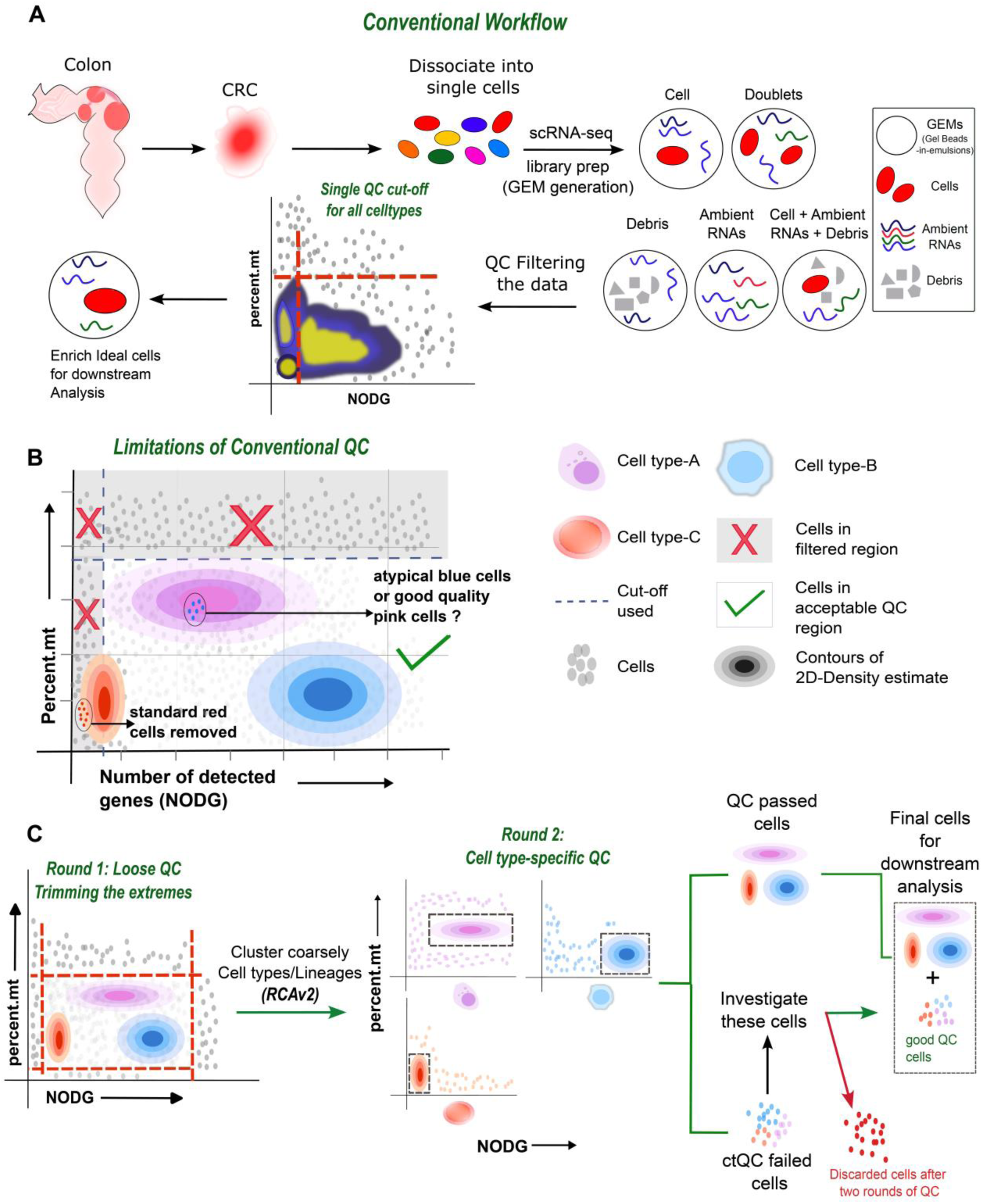
*ctQC* – data driven QC pipeline for scRNA-seq data. (**A**) Schematic depicts the workflow to obtain single cell expression data from primary tissues (CRC) and how conventional QC (hard thresholding) cutoffs are used to filter low-quality droplets. (**B**) Limitation of the most widely used data-agnostic quality control framework. Different density distribution as contours could be attributed to cell type specific distribution in QC metric space. (**C**) Illustration of Cell type-specific QC (*ctQC*) framework, which defines variable QC cut-offs (strict) in a cell type-specific manner.

Most commonly, cell QC involves imposing fixed cutoffs on key QC parameters such as number of detected genes (NODG), number of detected unique molecular identifiers (nUMI), and percentage of reads mapping to mitochondrial genes (percent.mt). For example, cell barcodes with fewer than 200 genes may be annotated as empty droplets and discarded (NODG<=200), and often an upper cutoff is also imposed on NODG to remove potential doublets [17]. Similarly, cells with percent.mt>30% may be discarded, since a high mitochondrial gene fraction may indicate low membrane integrity and low viability [18]. We refer to this QC strategy of using fixed, dataset-agnostic and cell type-agnostic QC cutoffs as conventional QC (Figure 1A, B). One key limitation of conventional QC is that a single cutoff may not be applicable to all datasets and all cell types since NODG and other QC parameters strongly depend on sample type, sample quality, cell type, reagents, and experimental protocol [19–22].

Due to the above limitations of conventional scRNA-seq data QC, newer algorithms use data-driven cutoffs. For example, *miQC* fits a two-component gaussian mixture model to percent.mt, conditioned on NODG: one component represents high-quality cells and the other low-quality [23]. It then discards cells based on posterior probability of belonging to the latter. One limitation of this method is that it assumes that all cell types have the same QC parameter distribution. To address this effect, *sctk* clusters cells in the space of QC parameters to define cell types based on data quality [24]. Then, for each such cluster, it fits a one-dimensional gaussian mixture model to the distribution of each QC parameter to define the corresponding high and low cutoffs. A potential limitation of this approach is that low-quality cells may not create discrete clusters in QC space. Rather, the distribution of cells in QC space could be continuous. Moreover, low-quality cells of one cell type may have the same QC metrics as high-quality cells of another type. *ddqc* performs high-resolution cell clustering in transcriptome space (res=1.3) to define provisional cell types. Then, for each cell type, a data-driven cutoff is set individually for each QC parameter based on standard criteria for outlier removal. The design philosophy of this algorithm is to discard as few cells as possible, and thus retain more cells than conventional QC [25], which could potentially result in retention of low-quality cells and debris. Similarly, *miQC* and *sctk* are tuned to retain approximately as many cells as conventional QC, and thus these methods may also retain debris and damaged cells.

We investigated the possibility that strict, cell type-specific QC (*ctQC*) could reduce data-quality artifacts in cellular-resolution transcriptomic data. To benchmark *ctQC*, we performed scRNA-seq on colorectal cancer (CRC) tumour samples kept on ice for varying time intervals to simulate sample deterioration after surgery. Cells retained by conventional QC but discarded by *ctQC* showed ambient RNA expression and ectopic expression of lineage markers. Relative to conventional QC and other data-driven methods, *ctQC* improved cell type separation in downstream cell clustering. Increasing the strictness of existing data-driven QC methods improved cluster separation, though *ctQC* remained the best performer. Analysis of published scRNA-seq data from fresh vs cryo-preserved human spleen yielded similar results. Relative to conventional QC, *ctQC* more clearly resolved patient-specific malignant cell states and CRC subtypes in an independent tumour scRNA-seq dataset [26]. Lastly, we found that *ctQC* was also applicable to spatial transcriptomes: the method improved accuracy in detecting anatomically localized cell types in Slide-seq data from human hippocampus [27]. To improve usability, we provide an R-shiny software package with a graphical user interface for performing *ctQC* (https://github.com/prabhakarlab/ctQC). Overall, our results demonstrate that strict, cell type-specific QC improves scRNA-seq and spatial transcriptomics data quality and thus improves our ability to resolve cell populations, characterize inter-patient heterogeneity and detect cell type-specific markers.

## Results

### Comparing *ctQC* to conventional QC

Although conventional single cell QC successfully removes empty droplets, we hypothesized that it may not completely remove debris and damaged cells, particularly when there is a delay between sample collection and single cell library generation (e.g. surgical or post-mortem tissues, banked samples or samples that require transport to a different site). To test this hypothesis, we generated scRNA-seq data from four aliquots of the same colorectal tumour sample, processed 0, 2, 4 and 6 hours after delivery (31,619 total cells, Fig.2A and Fig.S1A). For QC, we first tested the widely used strategy of imposing fixed cutoffs [13, 14, 17] on the number of detected genes (nFeature>=200 [28–32]) and the proportion of mitochondrial reads (percent.mt<=30 [28, 31, 33–35]). In addition, we used DoubletFinder to remove potential cell doublets [36]. We then examined the QC parameter distributions of the 23,589 remaining cells as a function of time. In contrast to 0-2h, the samples processed with a delay of 4-6h showed a marked increase in low-quality cells, i.e. cells with lower nFeature and higher percent.mt (Fig.2B). In particular, the QC distribution of the 4-6hr datasets showed two new low-quality modes with nFeature=500 and percent.mt=2% and 13% (Fig.2B, C). Since the four tissue aliquots were morphologically similar and derived from the same tumour sector, we did not expect major biological differences in their cell composition. Rather, we hypothesized that the low-quality QC modes that appeared at 4-6h were likely to represent damaged cells or debris from tissue degradation. Importantly, these putative low-quality cells were retained by conventional QC.

**Figure 2:**
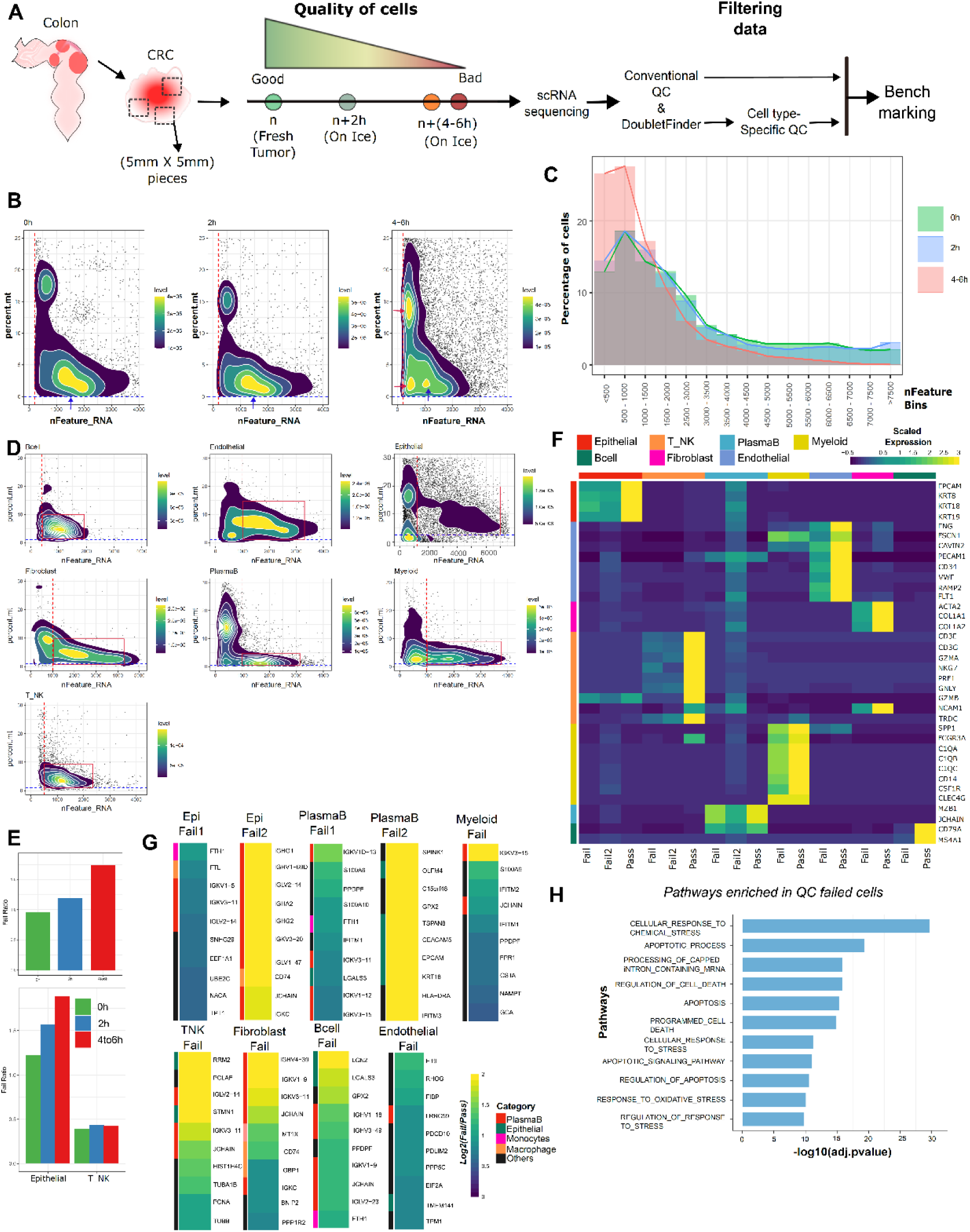
Conventional QC retains low-quality cells. (**A**) Schematic depicts experimental design and analysis strategy used in this study to benchmark *ctQC* pipeline. We collected samples from CRC patients and processed at different timepoints and later subjected to scRNA-seq. (**B**) QC metrics Scatter plot of all cells post conventional QC across different timepoints. Red and blue arrow indicates the mode of the distribution. (**C**) Binned histogram of nFeature_RNA across timepoint sample. (**D**) QC metrics scatter plot for broad individual cell types annotated by RCAv2 across all samples. (**E**) Failing odds ratio plotted across timepoints (top) and for individual cell types (bottom). (**F**) Heatmap shows expression levels of major cell type markers for cells that passed and failed *ctQC* cutoffs. (**G**) Top DE genes expressed in the QC failed cells compared to passed cells are plotted as heatmap. Genes are categorized to certain cell types based on their prominent expression. (**H**) Bar plots shows enriched pathways in QC failed cells derived using GSEA.

If the low-quality QC modes represented damaged cells, the most direct remedy would be to discard them by using stricter QC cutoffs. However, in doing so, one must take care to avoid discarding intact cells with intrinsically low nFeature (e.g. intact lymphoid cells). We therefore explored the use of cell type-specific QC (*ctQC*) cut-offs (Fig.1C): 1) apply conventional (loose) QC cutoffs; 2) cluster cells coarsely to define “QC cell types”; 3) apply stricter, lineage-specific cut-offs to each QC cell type; 4) re-cluster the remaining cells at the desired resolution. Coarse clustering was implemented using RCAv2, which clusters cells robustly in the presence of batch effects and technical variability [37]. RCAv2 identified 7 major cell types in the CRC scRNA-seq dataset (Fig.S1B). For each major cell type, the cut-offs for nFeature and percent.mt in Step 3 were defined based on the corresponding distribution of cells in two-dimensional QC space (Fig.2D, Methods), resulting in retention of 9,518 high-quality cells after the second round of QC.

In total, 14,071 cells retained by conventional QC were discarded in the second round based on strict cell type-specific cut-offs. We asked whether these 14,071 cells represented genuine cell types that should have been retained. We first examined the proportion of discarded cells as a function of processing delay and found that, as expected given their lower sample quality, the 4-6h samples had a higher overall discard rate [62.5% cells] (Fig.2E, Fig.S2B). This effect was stronger in epithelial cells [Failing odds – 0h: 1.22 Vs 4-6h: 1.9], which are known to be vulnerable to tissue dissociation and single cell processing [38], than in the more robust T cells [Failing ratio - 0h: 0.39 Vs 4-6h: 0.42] (Fig.2E, Fig.S2C). These results are consistent with the expectation that the 14,071 additional cells discarded by *ctQC* were of low quality.

To further investigate the nature of the 14,071 cells discarded by *ctQC*, we examined the expression of known markers for the 7 cell populations [39]. For each population, cells discarded by *ctQC* were labelled as “fail” and the rest as “pass.” Since epithelial and plasmaB cells included two distinct QC populations of failed cells (Fig. 2D), we further sub-categorised these cells as “fail1” and “fail2” (Fig.S2A). In all 7 lineages, discarded cells showed weaker expression of the respective marker genes, relative to pass cells (Fig. 2F). In particular, the fail2 epithelial and fail2 plasmaB subpopulations showed illegitimate expression of markers of other lineages (Fig. 2F, G). In summary, the cells discarded by *ctQC* showed relatively weak and inconsistent expression of lineage markers, indicating that their identity was less well defined, even at the lineage level.

Next, we directly evaluated the up-regulated genes in these *ctQC*-failed cells. For each lineage, we performed differential expression (DE) analysis to identify genes with higher expression in fail cells (Fig.S2A), compared to the corresponding pass cells (Methods). Interestingly, for each lineage, the top markers of fail cells coincided with markers of other lineages (Fig.2G). For instance, the strongest markers of fail epithelial cells were markers of the plasmaB cell type. Similarly top markers of fail plasmaB cells included epithelial and monocyte markers (Fig.2G), suggesting enrichment of ambient RNAs in these *ctQC*-failed cells. We then pooled all the up-regulated genes in *ctQC*-failed cells across the 7 lineages to identify the underlying biological functions and pathways. Gene set enrichment analysis (GSEA) identified enrichment of genes related to stress and apoptosis, again suggesting that the *ctQC*-fail population might be in a non-native state compared to pass cells (Fig.2H). Again, these results suggest that *ctQC* discards low quality cells retained by conventional QC.

The process of encapsulation in droplets can include ambient RNA (“soup”) in addition to intact/broken cells (Fig.1A). We used SoupX [10] to identify the 20 most highly expressed genes in ambient RNA in our dataset. Based on these 20 genes we used UCell [40] to calculate a module score for each cell, which we defined as the ambient RNA score. We noticed that the cells with elevated ambient RNA score overlapped in gene expression space with the 14,071 cells discarded by *ctQC* (Fig.3A). For example, epithelial and plasmaB cells with higher ambient RNA score were retained by conventional QC but discarded by *ctQC* (Fig.3A). A similar trend was observed for the other 5 cell types (Fig.S3B, C). Moreover, the ambient-high cells discarded by *ctQC* derived overwhelmingly from the samples processed after 4-6h (Fig.S3A). Lastly, *ctQC*-fail cells were transcriptionally distant from each other and from pass cells, i.e. the failed cells did not form coherent clusters (Fig.S4A). In contrast, pass cells showed strong cell-cell similarity (Fig.S4A). In summary, the 14,071 cells discarded by *ctQC* were associated with delayed sample processing, exhibited damage signatures, showed inconsistent expression of the expected lineage markers and high soup gene expression, and appeared to be transcriptomic outliers. We therefore concluded that these cells represented technical artifacts of sample processing rather than biologically distinct cell states.

**Figure 3:**
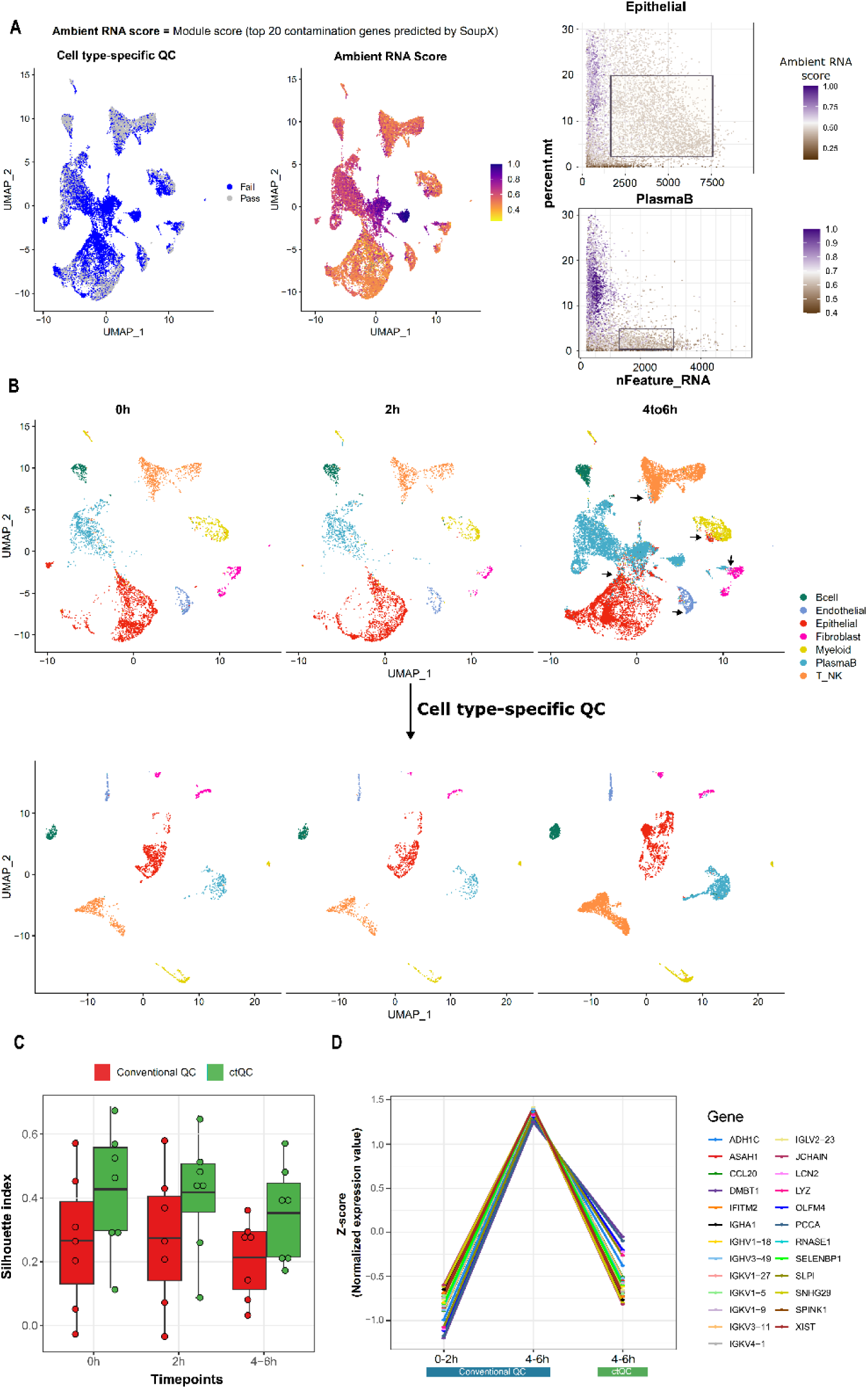
*ctQC* improves downstream analysis by removing debris. (**A**) UMAP coloured by ambient RNA score calculated from SoupX genes and by status of the cells whether they have met the ctQC cutoff criteria. Failed cells are highlighted in blue colour. Right, QC scatter plot for epithelial & plasmaB cells coloured by ambient RNA score and box defines the used nFeature_RNA and percent.mt cutoff. (**B**) Comparing the cell clustering in UMAP space post conventional and cell type-specific QC filtering. UMAP is split by timepoints. Black arrows indicate the regions where cells of two different types are intermingled in UMAP space. (**C**) Silhouette index derived based on broad cell labels are plotted across time points post conventional and *ctQC*. Black line in boxplot represents mean of the distribution. (**D**) Line plot depicts the scaled expression of top up-regulated genes in 4-6h epithelial cells compared to 0-2h epithelial cells.

Next, we asked if removal of low-quality cells could improve the quality of downstream cell clustering. To test this hypothesis, we clustered cells from the above dataset before and after *ctQC*. For a side-by-side comparison we used the same number of variable genes and principal components and adjusted the clustering resolution to equalize the number of clusters (Fig.S4B). Major lineages were more clearly separated after *ctQC*, both visually (Fig.S4C) and quantitatively in terms of silhouette index (Fig.S4D). Analysis of cells by timepoint showed that, with conventional QC, cell type separation worsened with increasing processing delay (Fig.3B, C). In contrast, after *ctQC*, cell type separation remained robust even with 4-6h of processing delay (Fig.3B, C). These results suggests that removal of low-quality cells from *ctQC* can improve cell type separation in downstream clustering analysis, particularly when samples are partially degraded.

To further explore robustness of conventional QC to technical factors, we identified genes differentially expressed between the 0-2h and 4-6h epithelial cells. Most strongly upregulated genes in 4-6h epithelial cells (highest fold-change) were a mix of plasmaB markers and ambient RNA genes predicted by soupX or cellbender, suggesting a relative increase in ambient RNAs due to sample degradation (Fig.3D, Fig.S4E). These genes showed significantly lower expression after *ctQC* (Fig. 3D, Fig.S4E). This result suggests that low-quality cells retained by conventional QC could bias gene expression estimates and potentially introduce artifacts in marker identification and other downstream analyses.

### Comparing *ctQC* to data-driven QC methods

We used the above CRC dataset to assess the performance of three previously developed adaptive, data-driven QC methods, *ddqc* [25], *miQC* [23] and *sctk* [24], as well as Cellbender, a clustering algorithm that adopts a distinct QC strategy based on ambient RNA estimation [16]. We used these methods at default settings to filter out low-quality cells and then discarded potential doublets using DoubletFinder [36]. Post-filtering, these methods retained 27,594, 25,802, 23,845 and 26,921 cells respectively (Fig.4A). Note that all four methods retained more cells than conventional QC (N=23,589). We used RCA2 to cluster these cells by lineage into 7 major cell types. Upon visual inspection of UMAP embeddings, we observed no clear qualitative improvement in cluster separation after QC by these methods (Fig.4A). For example, epithelial and plasmaB cells remained poorly separated and all lineages included subset of cells that were juxtaposed with a different lineage. This was particularly true of cells from the 4-6h samples, perhaps due to retention of debris and low-quality cells by these algorithms (Fig.S5A). Consistently, these four QC methods provided no discernible quantitative improvement in cell type separation (SI) relative to conventional QC (Fig.S6A). In fact, the SI of *ddqc*, *miQC* and *sctk* was even lower than that of conventional QC.

**Figure 4.**
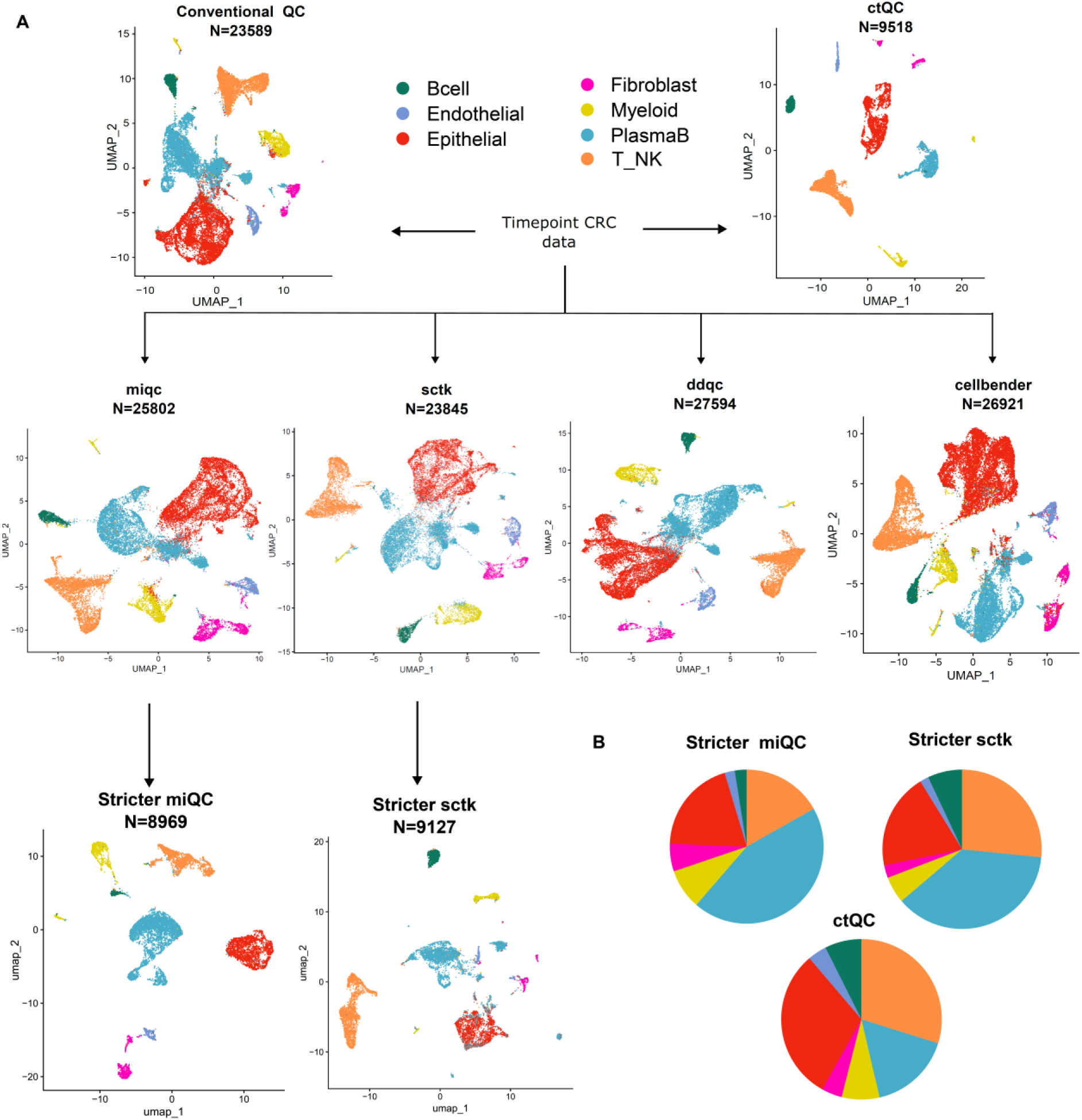
Benchmarking *ctQC* performance against existing QC pipelines. (**A**) Comparing cell clustering result of CRC timepoint dataset using three different QC pipelines and cellbender. All the programs were run using the default parameters and for *miQC, sctk* we also used stringent QC cutoffs. (**B**) Proportion of broad cell types post *ctQC* and stricter *miQC, sctk* represented as pie chart.

Since *ddqc* retained the largest number of cells, we investigated the 4,005 cells marked as high-quality by this method but discarded by conventional QC (ddqc_only). We compared these cells to those retained by conventional QC but discarded by *ctQC* (conv_only, 14,071 cells) and to cells retained by *ctQC* (ctqc, 9,518 cells). ddqc_only cells showed similar or weaker lineage marker expression relative to conv_only, while ctqc cells showed the strongest marker expression (Fig.S6B). Consistently, ddqc_only cells had relatively high percent.mt and lower NODG, suggesting inclusion of debris and low-quality cells (Fig.S6B, top red arrows). Though this trend was also observed in other cell types, it was most prominent in epithelial and plasmaB cells, which tend to be more vulnerable to degradation during sample processing (Fig.S6B). Thus, although both *ddqc* and *ctQC* apply cell type-specific cutoffs, the former may include marginal cells that weaken cluster separation in downstream analysis.

*ctQC* differs from conventional QC and existing data-driven methods in that it applies cutoff that are both cell type-specific and strict. To separate these two factors, we increased the strictness of *miQC* and *sctk* while retaining their cell type agnostic approach (it was not obvious to adjust the strictness of the other two methods). *miQC*_strict retained approximately the same number of cells (8,969) as *ctQC* (9,518), mostly by imposing, in effect, a stricter mitochondrial read threshold of 3.5% (Fig.S5B). This substantially improved cell type separation, though not as much as *ctQC* (Fig.4A, and S6A). Moreover, due to the strict mitochondrial cutoff very few cells with intrinsically higher mitochondrial percentage (B, Endothelial) were retained whereas plasmaB cells are overrepresented (Fig.4B pie). This overrepresentation suggest that stricter cutoffs retained low-quality plasmaB cells (Fig.S5D), given that plasmaB cells inherently exhibit low-NODG (Fig.2D). *Sctk*_strict also retained a similar number of cells (9,127) and again showed substantially better cluster separation that sctk but lower that *ctQC* (Fig.S6A). These results suggest that using stricter QC cutoffs can improve downstream clustering, and applying strict cutoffs in a cell type-specific manner can provide further improvements.

Overall, the above analyses of CRC time course data suggest that removal of debris and low-quality cells using strict, cell type-specific QC cutoffs can improve downstream clustering, marker identification. Thus, existing QC approaches, which primarily remove empty droplets and ignore cell type distinctions, may not be sufficient.

### *ctQC* resolves patient-specific cell states

Next, we examined a tumor scRNA-seq dataset we recently generated, based on which we defined two major malignant cell sub-types in CRC (iCMS2, iCMS3) [26]. This dataset included 46,055 epithelial cells post conventional QC, of which 13,912 were retained by *ctQC* (Fig.5A, B; Fig.S7A). For both QC methods we clustered the retained cells and evaluated patient specificity of the resulting clusters. Visual inspection of two-dimensional (UMAP) representations of cells in expression space suggested that *ctQC* was more successful than conventional QC in revealing inter-patient heterogeneity of malignant cells (patient-specificity of clusters; Fig.5A, B). To quantify this effect, we used the LISI metric for intermingling of cells from different patients in expression space [41] (Fig.5C) and further confirmed the same effect. Similarly, the silhouette index metric again showed that *ctQC* increased the patient specificity of the resulting embedding of cells in expression space (Fig.5D). Furthermore, iCMS2, iCMS3 and non-malignant epithelial cells were more well separated after *ctQC*, due to elimination of low-quality cells with ambiguous expression signatures (Fig. 5E, Fig.S7B, C). Thus, removal of low-quality cells by *ctQC* may improve our ability to resolve disease-related cell states and stratify patient by expression phenotype.

**Figure 5.**
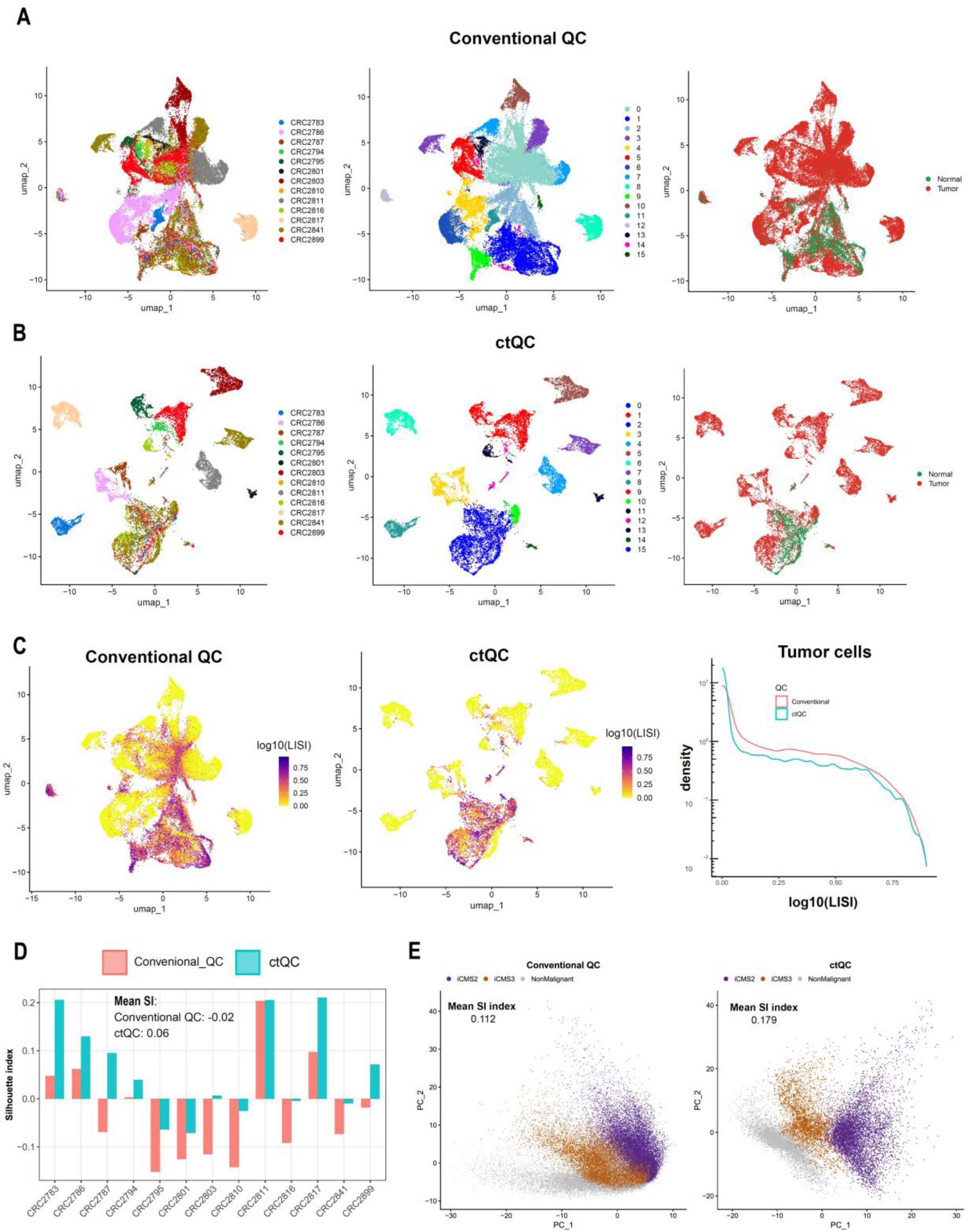
*ctQC* resolves patient specific clusters in tumor cells. **A, B**) Epithelial cells from Joanito etal was filtered using conventional and *ctQC*. UMAP are coloured by RNA clusters, patient id and tumour sample origin. **C**) UMAP coloured by log transformed Local Inverse Simpson index (LISI) score to look at patient intermixing. Density plot of log LISI score plotted for cells retained post conventional and *ctQC* (**D**) Silhouette index calculated patient-wise plotted as bar graph post conventional and *ctQC*. (**E**) Tumour cell stratified by molecular subtypes iCMS2, iCMS3 in principal component space.

### *ctQC* eliminates artifactual clusters in PBMC data

Next, we investigated the effect of *ctQC* on longitudinal PBMC scRNA-seq data from 33 COVID-19 donors and 14 healthy controls, comprising 246,964 adaptive and 125,117 innate cells [42]. First, we replicated the QC cutoffs used in the original study and defined the result as conventional QC. Multiple cell types were bimodally distributed QC metric space, suggesting the presence of distinct subpopulations (Fig.6A - D, Fig.S8A and Fig.S10A). In each case, one of the two modes showed low NODG and high percent.mt, and corresponded to a distinct cluster in transcriptome space (Figs. S9C, S11C). We hypothesized that these four clusters may represent technical artifacts, i.e. low-quality cells retained by conventional QC (Fig. S9A, Fig.S11A). Indeed, the vast majority of genes specific to these clusters appeared to be either mitochondrial genes or nuclear-enriched enriched transcripts that often characterize cells that have lost cytoplasmic RNA due to compromised membrane integrity (Fig.6F). In one case, namely CD16hi NK cells, the low-NODG cluster appeared to form a continuum in expression space between NK cells and classical monocytes, suggesting these cells may represent doublets between the corresponding cell types (Fig. S10B, C). The significant downregulation of ribosomal genes in these cells further supports the hypothesis that these cells may be technically biased (Fig.S9E, S11E). Since we were not aware of any studies of healthy PBMCs that detected such clusters, we asked whether they could represent disease-specific cell states. However, we did not observe any clear relationship between these clusters and disease state (Fig.6E). In summary, the above results suggest that the novel low-NODG and high-percent.mt clusters consistently observed across cell types in this dataset may represent artifacts of compromised membrane integrity, rather than distinct, disease-specific biological states.

**Figure 6.**
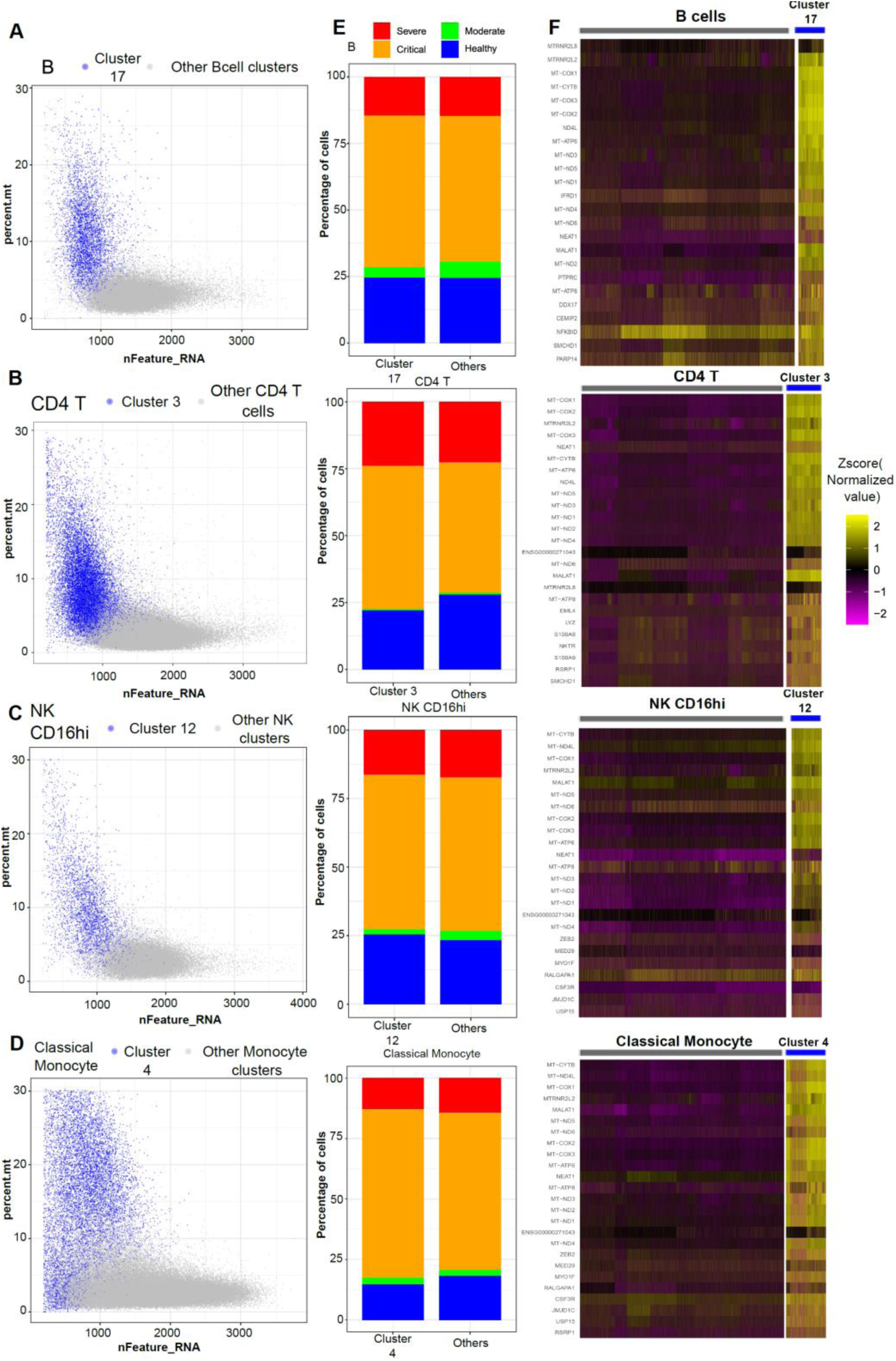
*ctQC* eliminates cells with compromised membrane integrity. (**A, B**) B and CD4 T cells from adaptive immune dataset are plotted in QC metric scatter plot. Low-quality cell cluster is highlighted in blue colour. (**C**, **D**) CD16hiNK and classical monocyte cells from innate immune dataset are visualized in QC metric scatter plot. Low-quality cell cluster is highlighted in blue colour. (**E**) Stacked bar plot representing proportion of cells stratified based on disease severity metric defined by the authors for low-quality cell cluster and other cells from the cell type. (**F**) Heatmap showing expression of top 25 marker genes for the low-quality cluster.

We investigated the ability of *ctQC* to address the above-described technical cell clusters. After *ctQC*, we retained 204,014 adaptive and 85,664 innate immune cells and clustered the cells once again following the method described the authors (Fig. S9A and Fig.S11A). Notably, the low-NODG technical clusters were no longer evident, and the intermediate cell state between NK cells and monocytes was no longer apparent (Fig.S8C and Fig.S10C). Moreover, cells that passed *ctQC* showed strong lineage marker expression relative to discarded failed cells (Fig.S9D and Fig.S11D). These results indicate that *ctQC* can enrich for intact PBMCs and reduce the prevalence of artefactual cell clusters that represent technical, rather than biological, effects.

### *ctQC* is applicable to data from preserved samples

Before single-cell library generation, primary tissues can be maintained for 1-3 days in a preservation medium to reduce sample degradation [43]. Since data generated in this manner may still contain an excess of low-quality cells, we hypothesized that *ctQC* could therefore provide improvements over conventional QC. We isolated primary tissue from a colorectal cancer patient and stored the samples in a preservation medium for up to 72h (Fig.S12A). Compared to 0h, we observed a reduction in the proportion of high-quality cells in data from the 24h sample,) suggesting a mild reduction in sample quality at this time point. This effect was more pronounced at 72h suggesting greater degradation of this sample (Fig.S12B). As above, we used RCAv2 to cluster cells by lineage and performed *ctQC*. Indeed, we found that *ctQC* improved cell lineage separation, both visually and by the quantitative SI metric (0.28 vs 0.36, Fig.S12D-F). We performed a similar analysis of scRNA-seq data from human spleen, generated fresh as well as after 72h of storage in a preservation medium [44]. For conventional QC, we applied the cut-offs used in the original study (nFeature >=300, percent.mt<=10%) and retained a total of 49,391 cells from the two timepoints. As expected, we observed lower cell quality metrics in the 72h sample (Fig.S13A), and *ctQC* discarded more cells in this sample (Fig.S13B). Again, *ctQC* improved cell lineage segregation in expression space (Fig.S13C), presumably by discarding low-quality cells with relatively weak expression of lineage markers (Fig.S13D). These results suggest that conventional QC retains low-quality cells in scRNA-seq data from preserved samples and that *ctQC* could therefore improve the quality of downstream clustering.

### *ctQC* is applicable to spatial RNA sequencing

To evaluate the applicability of *ctQC* to spatial RNAseq data we first examined slide-seq v2 data from mouse hippocampus [27], comprising spatially localized transcriptomes of 53,173 pucks. We retained these pucks based on author’s cutoff as conventional QC. For *ctQC* the pucks were grouped into cell types based on the authors’ annotations and QC cutoffs were applied in a cell type-specific manner resulting in retention of 27,152 pucks. The two post-QC datasets were clustered as in the original study and clusters were labelled based on the original annotations. We noticed that the dentate principal cell cluster based on conventional QC included a subpopulation of pucks with low NODG and high percent mito (Fig. 7B). These pucks, which were spuriously located outside the dentate gyrus, were discarded by *ctQC*. To quantify this observation, we defined two measures of spatial coherence of pucks within the same cluster: 1) proportion of neighbouring cells that have the same type as the index puck (neighbourhood homogeneity) and 2) distance to the closest puck of the same type (homotypic distance). Cells discarded by *ctQC* had relatively low neighbourhood homogeneity and high homotypic distance, indicating that *ctQC* discarded pucks with low spatial coherence that tended to lie outside the dentate gyrus (Fig. 7B). To extend this analysis, we examined Slide-seq v2 data from mouse cerebellum [45]. Again, we found that clusters based on conventional QC included mis-localized pucks that lay outside the expected anatomical regions for Bergmann glia and Purkinje neurons (Fig. 7C, D). In contrast, post-*ctQC* clusters for these two cell types showed greater spatial coherence and anatomical fidelity. These results suggest that *ctQC* can improve accuracy in detecting anatomically localized cell types in spatial RNA-seq data.

**Figure 7:**
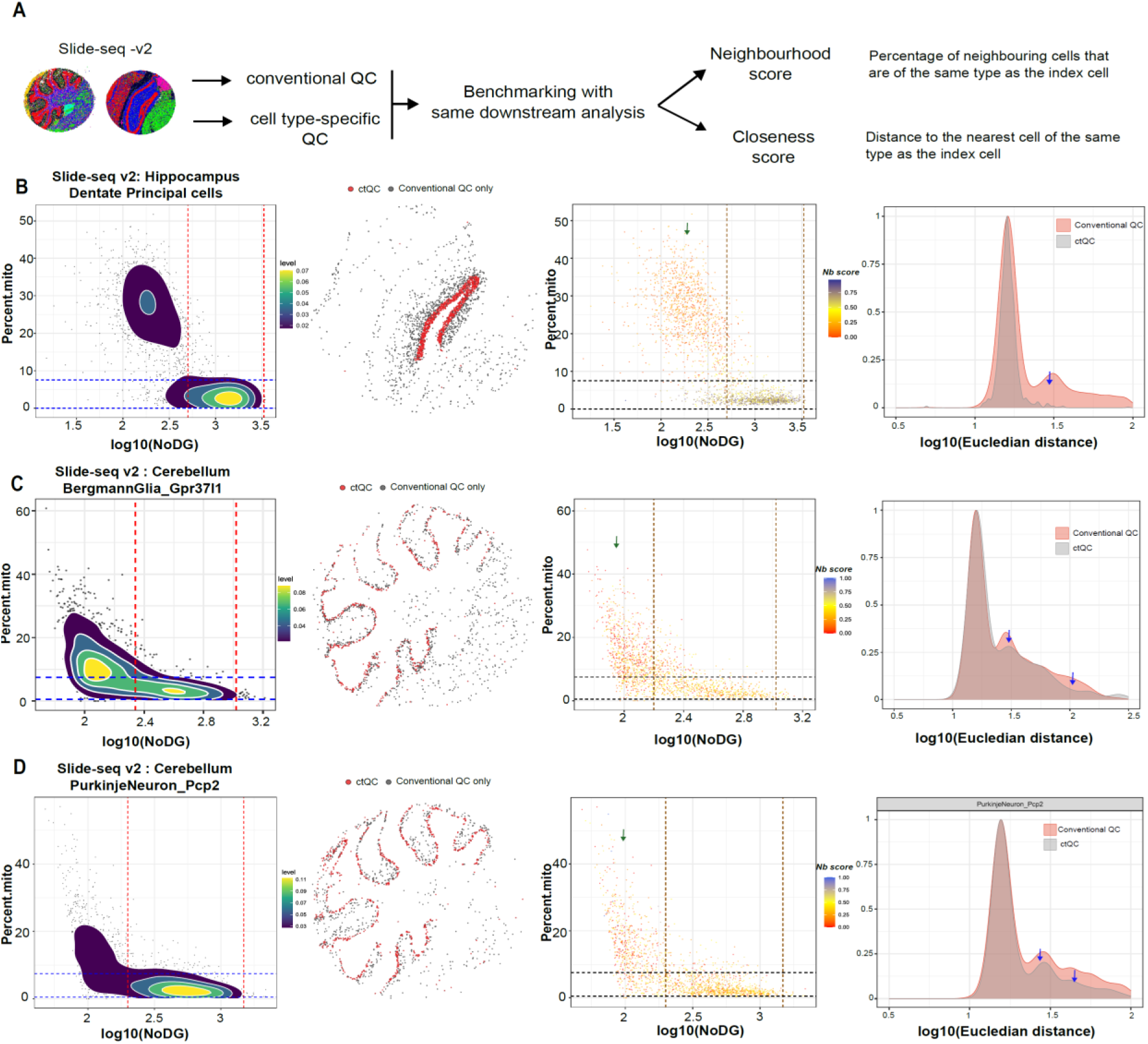
*ctQC* is applicable to sequencing based spatial transcriptome datasets. (**A**) Analysis strategy used to evaluate effect of *ctQC* on slide-seq data. (**B**) Dentate principal cells from hippocampus slide-seq dataset are plotted in QC metrics scatter plot. Red & blue lines indicate the *ctQC* cut-offs used. In the next plot, spatial distribution of pucks retained post *ctQC* are coloured in red while pucks retained post conventional QC is in grey. In next plot, calculated neighbourhood scores are overlaid in QC metric scatter plot. Finally, closeness score calculated post conventional and *ctQC* are plotted as density plot. (**C**, **D**) Similar to (**B**) for two cell types (Bergmann Glia and Purkinje neuron pcp2) from cerebellum slide-seq dataset are plotted.

## Discussion

Currently, scRNA-seq studies use conventional QC with relatively loose thresholds that efficiently remove empty droplets but may retain debris. Similarly, conventional QC can retain cells whose mitochondrial read percentage substantially exceeds the mode of the distribution (>2-fold or >3-fold). The objective of this approach is to increase statistical power by maximizing the size of the post-QC dataset. However, QC cutoffs cannot be loosened indefinitely, since data points of very low quality could introduce artifacts that obscure or distort the underlying biology. The optimal trade-off between dataset size and cell quality is thus a key consideration in single cell data analysis. Similarly, spatial omics data analysis also requires a balance between number of cells and cell quality.

We implemented strict QC cutoffs (*ctQC*) and found that clustering results improved by multiple metrics, including cell type separation, lineage marker specificity, spatial contiguity, and resolution of patient-specific cell states. Simultaneously, artefactual clusters consisting almost exclusively of low-quality cells were minimized. This trend was consistent across the single cell and spatial datasets we analyzed: CRC, human spleen, PBMC and mouse brain. The improvement from strict QC was potentially attributable to the fact that conventional QC retained an excess of cells in the “grey zone” that lies in between completely empty droplets and intact cells (Figs. 2, 3). As expected, these low-quality cells were more abundant in tumor samples that faced a longer delay in processing (Figs. 2-4). Overall, our results suggest that conventional scRNA-seq and spatial QC cutoffs may be too lenient, particularly in the case of primary-tissue datasets containing a substantial proportion of damaged cells.

One straightforward remedy would be to increase the stringency of existing QC methods, both conventional and data-driven. Indeed, we found that this strategy could improve downstream cell clustering, but only to a point (Fig.4). A key limitation of this strategy is that QC metrics such as NODG and percent.mt strongly depend on cell type (Fig.2D, S8A, S10A, S14A and S15A). Consequently, a one-size-fits-all QC cutoff can be too strict for one cell type and yet too lenient for another. Our *ctQC* approach addresses this problem by setting QC thresholds in a cell type-specific manner, so as to maintain a consistent degree of stringency across cell types.

We note that the QC parameter distribution of a cell type can vary across studies, due to differences in the assay (reagent kit, protocol) as well as differences in sample quality. Consequently, we have adopted a data-driven approach for determining QC thresholds based on the mode of the two-dimensional distribution. Some existing methods are also data-driven, for example *ddqc* and *sctk*. However, these methods did not improve cell clustering relative to conventional QC, perhaps because they set lenient cutoffs that retained even more cells than the latter.

Strict QC raises the concern that one could in the worst-case discard genuine cell types, for example cells that are metabolically active and therefore express mitochondrial genes at a higher level[19, 46, 47]. Similarly, cell types such as granulocytes, which have intrinsically low NODG in scRNA-seq data, could potentially be discarded. In the datasets we examined, we found no evidence that the discarded cells represented novel cell types or distinct biological states. Perhaps this is because our cutoffs were set in a cell type-specific manner, which accommodates intrinsic biological differences between cell types. Nevertheless, we recommend that discarded cells be examined carefully for lineage marker expression and DE genes, to confirm that they are indeed technical rather than biological.

A limitation of our approach is that it examines only two QC parameters: NODG and percent.mt. One extension would therefore be to also incorporate the number of detected RNA molecules (nUMI). However, in practice nUMI is highly correlated with NODG within any given cell type. Thus, inclusion of nUMI is unlikely to substantially affect the results of *ctQC*. Nevertheless, it may be beneficial to explore extensions of *ctQC* that perform strict, cell type-specific and data-driven QC in a higher-dimensional space that incorporates a larger set of QC metrics. We note also that *ctQC* may not be needed when the vast majority of cells are of high quality. In such cases, the number of discarded cells may be too small to make an appreciable difference in downstream analyses.

To simplify the process of performing *ctQC*, we have developed an interactive R-shiny package that accommodates both scRNA-seq and sequencing based spatial omics data (https://github.com/prabhakarlab/ctQC). The *ctQC* package implements the QC workflow used in this study (Fig. 1C), including exploratory analysis of discarded cells to evaluate the possibility that they might represent genuine biological types or states. To guide selection of subpopulations, the package fits Gaussian mixture models (GMMs) to the distribution of cells in QC space for each cell type.

Overall, our results suggest that single cell and spatial QC methods may benefit from combining three attributes: 1) data driven, 2) cell type-specific and 3) strict. QC algorithms that incorporate these features could improve our ability to extract biological signals from single cell and spatial omics data.

## Material and Methods

### Sample collection

CRC timepoint and hypothermosol datasets generated in this study are from a single colorectal cancer patient (Patient id: CRC3526). The samples collected from this patient were in accordance with ethical guidelines [IRB: 2018/2795 for Circulating Markers and Molecular Biology in Colorectal Cancer or Colorectal Neoplasms] and the patient provided written, informed consent. After resection, tumor colon tissues were collected and cut into multiple smaller pieces (approximately 5X5mm) for downstream processing. Each tissue aliquot was processed as per our experimental design shown in Fig.2A and Fig.S12A. The control sample (0h) was immediately transferred for tissue dissociation, while 2h, 4h and 6h samples were kept on ice for subsequent time point processing, and 24h and 72h samples were stored in hypothermosol for later dissociation. Irrespective of the timepoint, all sample processing and library preparation procedures were kept uniform throughout.

### Library preparation

The transportation medium (hypothermosol) was decanted out and tissue specimens were placed in petri dishes, weighed, and kept on ice for dissociation. Tissues were then subjected to fine mincing in 5 ml of RPMI solution with 10% FBS (Gibco, Life Technologies) and 1% Pen/Strep (Gibco, Life Technologies) using sterile scalpel blades to make a fine slurry. The minced tissue was resuspended in pre-warmed dissociation buffer 2X, comprising Collagenase-P (4 mg ml−1, Roche) and DNase-I (0.4 mg ml−1, Roche) in a total of 5 ml of RPMI medium containing 10% FBS (Gibco, Life Technologies) and 1% Pen/Strep (Gibco, Life Technologies). The tissue suspension was enzymatically digested using two rounds of shaking incubation at 37 degrees for 6 min each with 1 min of vigorous vortexing in between. The enzymatic digestion step was then followed by one more round of physical homogenization by pipetting the tissue suspension through 10-ml and 5-ml pipette bores for at least 1 min. The resulting suspension was washed with an ice-cold Dissociation Buffer (1% BSA / DPBS / 2mM EDTA) and the contents were passed through a 70-µm strainer to remove undissociated tissue. The filtrate (single-cell suspension) was centrifuged at 500g for 5 min at 4 °C and the supernatant was decanted. To remove red blood cells, the cell pellet was treated with ACK lysis buffer (Gibco, Life Technologies) for 5 min on ice. Following red blood cell lysis, the sample was washed again with ice-cold Dissociation Buffer, re-filtered through a 40-µm strainer and centrifuged at 300g for 5 min at 4 °C. The supernatant was decanted without disturbing the pellet and resuspended in 3–5 ml of RPMI solution (depending upon pellet size). 10 µl of this cell suspension was mixed with 10 µl of tryphan blue dye to perform cell counting and viability using an automated cell counter (Luna).

### 10X library Sequencing

Six processed samples of varying time points (0h,2h,4h,6h,24h and 72h) right after dissociation were loaded into the Chromium controller system (10X Genomics) with a target of 7,000 cells per well. Barcoded sequencing libraries were generated using the Chromium Single Cell 5′ v1.1 dual index reagent Kit. All libraries were sequenced on an Illumina NovoSeq S4 flowcell until sufficient saturation was reached.

### Analysis of CRC timepoint data by conventional QC

Raw scRNA-seq reads were assigned to cells (barcoded droplets) using CellRanger v.6.0.2 (reference - gex-GRCh38-2020-A) [48] to generate the raw expression matrix of unique molecular identifier counts (UMI counts; indicative of number of unique RNA molecules detected) for each gene in each cell (droplet). Droplets with number of detected genes (NODG) < 200 were discarded as empty droplets. After discarding empty droplets, we retained 5,435, 4,152, 9,178 and 12,854 cells from the 0h, 2h, 4h and 6h samples respectively (Fig.S1A). We pooled cells from the four timepoints and performed pre-processing, broad lineages identifications and downstream analysis. We then performed data-agnostic conventional QC by setting cutoffs as: nFeature_RNA <= 200; percent.mt >= 30 and nCount_RNA <= 500. Post conventional QC, we retained 24,978 cells across all four timepoints. We then used DoubletFinder v.2.0.3 [36] for each sample (timepoint) separately to remove putative doublets from the dataset. Here, we followed the suggested workflow, with doublet rates of 0.8% per 1,000 recovered cells (following 10X Genomics protocol). We removed 1,389 cells as doublets and finally used 23,589 cells for all the downstream analysis.

The data was then logNormalized with scaling factor = 10,000 using Seurat v4.3.0 [14]. To identify broad cellular lineages, we projected the log normalized single cell transcriptome data onto the Global panel of RCAv2 (using Pearson’s correlation) [37]. 23,589 cells were clustered at lower resolution using RCAv2 with default parameters and the function ‘estimateCellTypeFromProjectionPerCluster’ was used to broadly annotate cell types for each identified cluster. We identified 7 major cell types from the data: Epithelial, B, endothelial, plasmaB, T_NK, myeloid and fibroblast. Fig.S1B shows the heatmap of projection of cell types to clusters as well as the table summarising the number of cells per cell type. We used Seurat v4.3.0 to perform standard downstream analysis (normalization, finding highly variable genes (n= 2000), data scaling, PCA, clustering and visualization; Fig.S4B).

### Cell type-specific QC (*ctQC*)

*ctQC* is a two-step data-driven QC pipeline where cells are subjected to first round of lenient QC followed by second round of strict cell type-specific thresholding (Fig.1C). The first round of QC cutoffs is to remove obvious low-quality cells and empty droplets from the data. The second round of QC is strict and cell type-specific, which maintains uniform strictness across cell types within a dataset. To perform the second round of QC we need broad cell type annotation and we used RCAv2 with default parameters to cluster the cells in lower resolution and to annotate the clusters. ctQC thresholding is jointly decided based on library complexity (NODG) and the percentage of mitochondrial reads (as a measure of viability/integrity [18]). For each cell type, this QC metric scatter plot (NODG Vs percent.mt) is generated using ggplot2 [49] and the density distribution of points (cells) is overlayed using stat_density_2d function (https://ggplot2.tidyverse.org/reference/geom_density_2d.html), such that the calculated 2D kernel density estimation is plotted as contours in the scatter plot. We use the density contours to identify the mode(s) in the given distribution. We observed an inherent cell type or lineage specific QC distribution in the 2D scatter plot highlighting the necessity of tailor-made cutoffs in a cell type specific manner to retain biologically meaningful cells (Fig.2D).

For every broad cell type, we defined upper & lower thresholds for NoDG and percent.mt as being within X deviations from the 2D-mode of the distribution. Several factors such as experimental design, sample processing, tissue type and cell lineage can influence the choice of strictness. By default, we use 2.5X deviations from the mode of the QC metrics distribution as cutoff for all cell types in a dataset. In our experience, 2-2.5X standard deviations from the mode worked well for all datasets analysed in this study. For instance, we used 2.5X for mouse brain data, 2X for Covid PBMC data, 2X for spleen data. For our CRC time point data (data generated in Fig.2A) we modified the cutoffs for a few selected cell types to ensure we retained high quality cells.

### *ctQC* for CRC timepoint dataset

For our CRC time point data, we used the same cutoffs used for conventional QC (NODG >= 200; percent.mt <= 30) as the first-round cutoff (loose cutoff), so that the broad cell type labels predicted post conventional QC can be used to perform *ctQC* (Fig.S1B). For the second round of QC, we plotted the QC metric scatter plot for individual cell types with stat_density_2d (Fig.2D). Since the dataset had samples that are deteriorating to different extents, we modified the cutoffs for certain cell types from a factor of 2X deviations from the mode. In case of epithelial cells, the upper percent.mt & NODG thresholds were retained at 2X deviations from mode (mito <= 20 & NODG <= 7000) while the lower percent.mt & NODG thresholds were adjusted as appropriate (mito >= 2.5 instead of 5 & NODG >= 1250 instead of 1750). We found that these cutoffs removed most of the low-quality cells and retained a higher proportion of good quality cells. Other tested cutoffs for epithelial cells either retained more low-quality cells or discarded more good quality cells. Similarly, we had to modify the upper or lower cutoffs for few other cell types in the dataset. We further performed an in-depth investigation to examine cells exclusively discarded by *ctQC* (fail categories in Fig.S2A and Fig.2) are indeed low-quality (see description below).

### Investigating ctQC failed cells

We labelled the 14,071 cells that were discarded by *ctQC* as “fail” and further segregated these cells by their broad cell lineages for downstream analysis. PlasmaB and epithelial cells have two prominent percent.mt modes and are labelled as “fail1” and “fail2” (Fig.S2A). We did single cell DE analysis between the cells described in Fig.S2A and their corresponding *ctQC* ‘pass cells’ for each cell type using the FindMarker function from Seurat [14]. The top 10 up-regulated genes in the “fail” category for every cell type were plotted as heatmaps using pheatmap (https://github.com/raivokolde/pheatmap, Fig.2G). We then pooled all the up-regulated DE genes (adj.pvalue < 0.01 & foldchange >= 1.5) across cell types and did enrichment analysis using GSEA [49]. We calculated cell-cell correlations between ctQC “passed” and “failed” cells for all cell lineages over the 0-2h and 4-6h timepoints. We calculated correlations using the “cor” function in R [50] and plotted the values using pheatmap (Fig.S4A).

### Ambient RNA analysis

We used SoupX [10] and cellbender [16] to predict the ambient RNA or genes that are enriched in soup from our CRC timepoint dataset. To get contamination profiles, SoupX uses raw and filtered cell ranger outputs to identify empty droplets. It ranks the genes in these empty droplets and identifies genes that are enriched in soup. To identify the cells which have enriched ambient RNA expression, we took the top 20 ambient RNA genes predicted by SoupX and calculated single cell gene signature scores using UCell [40]. This UCell score is plotted in Fig.3A and Fig.S3. We also used cellbender to identify systemic background noise in the data. We ran cellbender with default parameters (without the expected-cells parameter) using the cell ranger raw-feature matrix as input. The top 10 removed genes from cellbender were then used to calculate the ambient RNA proportion in the data. We merged the genes predicted from these two programs for Fig.S4E.

### Benchmarking analysis

The raw cell ranger output of CRC timepoint data (31,619 cells) was analysed using *miQC* [23], *sctk* [24] and *ddqc* [25]. *miQC*, utilizes a probabilistic framework model that jointly models NODG and percent.mt to eliminate low-quality cells. By default, *miQC* removes cells with 75% or greater posterior probability of being compromised. We retained 25,802 cells post *miQC* using the default cutoff; to increase the stringency of the program we modified the posterior probability to 0.01, which resulted in the retention of 8,969 cells (Fig.S5B). *sctk* clusters cells based on QC parameters (nUMI, NODG, percent mito, ribo, hb; instead of transcriptional variation) and uses gaussian mixture models to define high/low cutoffs for the individual QC parameters. We retained cells that passed both the “per_cell_qc” and “qc_cluster” calls. *Sctk* with default cutoffs retained 23,845 cells and with increased stringency (probability density function of mixture model threshold to 0.001) retained 9127 cells (Fig.S5B).

*ddqc* performs high-resolution clustering of data (post lenient cutoffs – NODG >100 & percent.mt <= 80) and defines cluster wise adaptive NODG, percent.mt and percent.ribo cut-offs (default cutoff is 2.5X of Median Absolute Deviation). *ddqc* with default cutoffs retained 27,594 cells. We grouped cells as “ddqc_only”, “conventional_only” based on the strategy explained in the diagram shown in Fig.S6B. Post grouping, we used Seurat’s AverageExpression function to get pseudobulk expression. Pseudobulk expression was then used to plot the heatmap. We used cellbender to annotate cells from our CRC timepoint dataset. As intended, the cellbender identified more cells (Cell ranger N: 31,619; Cellbender N: 41,698) from the data. However, ∼34% (N=14,002) of the annotated cells had NoDG<100 indicating that a huge proportion of low-quality droplets are annotated as cells. We used similar cutoffs to those in the cellbender paper [16] (nFeature_RNA <= 100, percent.mt > 75^th^ percentile and nCount_RNA > 85^th^ percentile are removed) in our dataset and retained 26,921 cells. Since all four methods retained more cells than conventional QC, we ran RCAv2 separately for each method to identify broad cellular lineages for these cells. Downstream analysis post QC for all the four methods was kept uniform (Fig.S4B). To calculate the overlap of retained cells across methods and strictness we plotted an upset plot (Fig.S5D) [50]. We plotted the overlap for two cell types (PlasmaB and T_NK) which showed proportion differences post strict QC cutoffs (Fig.4B). PlasmaB cells proportion differences in Fig.4B could be due to 1067 low-quality plasmaB cells discarded by ctQC were retained by all the methods (Fig.S5D).

### Calculating silhouette index

The silhouette index is a measure of how well each point fits within a given group compared to other groups, thereby indicating the quality of clustering. In particular, higher values imply better clustering. For all the datasets, post conventional and *ctQC* filtering, we did downstream analysis as shown in Fig.S4B and adjusted the resolution to get similar number of clusters. We then used the function SilhouetteBenchmarkPerCTPCA from RCAv2 [37] to calculate the silhouette index for all the datasets.

### Tumor cell analysis from CRC-SG1 cohort

From our previously published CRC single cell data [26] we extracted only the epithelial cells for this study. The CRC-SG1 cohort had a total of 79,978 epithelial cells; we used the cutoffs NODG >= 300; percent.mt <=30 as conventional QC and retained 46,055 cells. For *ctQC*, based on scatter plots of the QC metric, we used a cutoff of NODG >= 2200 & NODG <= 7500; percent.mt >= 3 & percent.mt <= 23 and retained 13,912 cells (Fig.S7A). Cells post conventional QC and *ctQC* were analysed similarly as shown in Fig.S4B. To calculate the intermixing of tumour cells between patients we calculated Local Inverse Simpson’s Index (LISI) scores using the R package LISI (https://github.com/immunogenomics/LISI) [41]. We log-transformed the LISI score and used it for further analysis. Tumor molecular classification (iCMS2, iCMS3) and tissue origin were obtained from the metadata provided from the original study.

### Covid PBMC analysis

We downloaded published covid PBMC data [42] where 264,964 adaptive and 125,117 innate immune cells were sequenced over the time course of COVID-19 infection. We used the cells retained by the authors as conventional QC, as well as their broad cell-type annotations, patient metadata, and downstream analysis pipeline. To mirror the original study, we analysed innate and adaptive immune cells separately. Most of the individual cell type QC metric plots (NODG Vs Mito) had a sub-population of cells in the low-NODG-high-mito region (Fig.S8A, S10A). We used *ctQC* cutoffs as 2X deviations from the mode of the distribution and retained 204,015 adaptive and 85,664 innate cells post *ctQC*. Most of the cells from these low-quality clusters were discarded by *ctQC* (Fig. S9B and S11B). We did single cell DE analysis for low-quality clusters (3,4,12 and 17) against other cells from that cell type using the Seurat FindMarkers function (Wilcoxon test). DE genes were sorted by adjusted-pvalue (*P* < 0.01) and fold change (>= 2 fold) values to identify the top 25 upregulated and downregulated genes in these cells (Fig.6F, S9E and S11E). Markers for these cell types were obtained from the original study (Fig.S9D, S11D). Heatmaps generated using R pheatmap package.

### CRC hypothermosol sample analysis

We generated scRNA-seq datasets from CRC tissue stored in hypothermosol for 24h and 72h as shown in Fig.S12A. We sequenced a total of 7730 and 7770 cells from the 24h and 72h samples respectively and used the 0h sample from our CRC timepoint data as control. Post conventional QC (NODG >= 200 & percent.mt <= 30) and doublet removal we retained 6018(24h) and 4476(72h) cells. In the 72h sample, the overall mode of the NODG distribution was below 1000 and we observed more cells with higher percent.mt indicating low-quality cells (Fig.S12B). We then used RCAv2 to identify broad cell types and performed *ctQC* similar to Fig.2D. For all cell types, based on the 2D mode of their distribution in the QC metric plot, we used a cutoff of 2X deviation from the mode. We retained 3,992(24h) and 2,270(72h) cells post *ctQC*. We used similar downstream pipeline as shown in Fig.S4B to analyse cells post conventional and *ctQC*. In UMAP space, we observed more bridging cells post conventional QC and less intermixing of cells post *ctQC* (Fig.S12C, F).

### Human spleen analysis

We downloaded published human spleen scRNA-seq data [44] where spleen samples were stored in preservative medium for 72h (Fig.S13A). We used the control 0h and 72h samples for the purpose of this study (0h – 18,024 & 72h – 31,367 cells). Similar to our CRC 72h hypothermosol sample, spleen 72h samples also had the mode for the NODG distribution below 1000 (Fig.S13A). We used the author-provided cell type annotations to perform *ctQC*. For all broad cell types we used 2X deviation from the mode to define *ctQC* cutoffs and retained 43,250 cells post *ctQC* (0h – 16,235 & 72h – 27,015 cells). *ctQC* discarded more cells from the 72h sample (Fig. S13B) and most of the discarded cells were bridging cells between different cell types. Post *ctQC* there was less intermixing of cells and cells discarded by *ctQC* had relatively lower expression of marker genes (Fig.S13D). Pseudobulk values were obtained using Seurat’s AggregateExpression function and used for plotting the heatmap (Fig.S13D).

### Slide-seq data analysis

We downloaded the slide-seq data for mouse hippocampus [27] and cerebellum [45] from the broad institute’s single-cell portal (https://singlecell.broadinstitute.org/single_cell). We had a total of 53,173 pucks from the hippocampus dataset and 27,196 pucks from the cerebellum dataset (RCTD annotated pucks [45]). We used author-provided puck labels as broad cell types to perform *ctQC*. The QC distribution for all cell types is shown in Fig.S14A and S15A. We used 2.5X deviations from the mode of the distribution as *ctQC* cutoff and retained 42,769 pucks from the hippocampus dataset and 18,849 pucks from the cerebellum dataset. *ctQC* cutoff boundaries for all cell types are shown in Fig.S14A and Fig.S15A. We used Seurat’s spatial module (https://satijalab.org/seurat/articles/spatial_vignette) to analyse the data post conventional and *ctQC*. We normalized the data using SCTransform [51] and used the same number of variable genes and principal components (PCs) for downstream analysis.

Neighbourhood and closeness scores were calculated for both slide-seq datasets. Both scores are defined in Fig.7A. We used customized perl scripts to calculate the scores and plotted them using the R package ggplot2 [49]. The closeness score was plotted as a density distribution comparing conventional and *ctQC* for all cell types (Fig.S14C and S15C). Spatial distribution of the pucks for different cell types post conventional and *ctQC* were plotted using Seurat’s SpatialDimPlot function (Fig.S16).

### *ctQC* shiny implementation

We have developed an R-shiny package (https://github.com/rstudio/shiny) that is easy to use and user-interactive. This package has three major steps as shown in Fig.1C that are implemented to perform *ctQC*. The dataset can be input using 3 different formats 1) as a seurat or scanpy object; 2) as a Gene X Cell matrix; or 3) as a .csv or .tsv file with NODG, percent.mt and broad cell type annotations. Grouping cells into broader lineages can be done using the package (if the input is of format type 1 or 2); or the user can input the broader group label (input format 3) based on which cell type-specific QC cutoffs can be decided. For every cell type, QC metric scatter plots with density contours are overlaid so that the user can pick cell type-specific thresholds. We have also implemented mixed gaussian models (based on NODG and percent.mt) for every cell type; users can pick a model (number of mixture components can be adjusted) that better fits the QC distribution. Cells that pass cell type-specific QC thresholds could be downloaded for every cell type. There is also an additional option to probe the quality of cells that are discarded by *ctQC*. For large datasets we recommend importing input data in the format 3 for faster results. Package can be accessed from the following github repository https://github.com/prabhakarlab/ctQC.

## Data Availability

CRC timepoint and hypothermosol datasets are deposited in European Genome-Phenome Archive (EGA) under the accession id EGAD50000000534. We downloaded the covid PBMC data for both adaptive & innate immune dataset as cellXGene matrix deposited from (https://cellxgene.cziscience.com/collections/ed9185e3-5b82-40c7-9824-b2141590c7f0). We downloaded the mouse brain slide-seq hippocampus & cerebellum data (pucks) from (https://singlecell.broadinstitute.org/single_cell). Spleen hypothermosol data was downloaded from tissue stability atlas (https://www.tissuestabilitycellatlas.org/). CRC-SG1 epithelial cell data is from a previously published study [26] from the lab (EGAD00001008555).

## Author contributions

S.P, I.B.T and V.L designed and conceived the project. V.L analysed all the data and interpreted the results with S.P; C.M., C.F. performed the surgery and isolated the tissue. L.W.Q. provide pathology support. M.K., P.N.V., D.C.N., N.G. processed the sample (primary tissue) and generated the scRNA-seq library. V.L and S.P wrote the manuscript with critical inputs from other authors. S.P and I.B.T supervised the study. All authors have read and approved the final contents in this manuscript.

## Acknowledgements

This work was supported by National Medical Research Association Clinical Scientist – Senior Investigator Award (grant no. MOH-CSASI22JUL-0003 (I.B.T)); the National Medical Research Council, Singapore (grant no. OFIRG21jun-0090 (S.P.)); the National research Foundation fellowship, Singapore supported P.N.V. (grant no. NRF-CRP29-2022-0005). We would like to thank Dr. Kian Hong Kock for inputs and discussions. Single cell sequencing was performed at GIS core facility - Spatial and Single Cell Genomics Platform (S2GP). The authors would also like to thank Dr. Jonathan Aow, Dr. Reema Baskar, Dr. Jagadish Shankaran and Dr. Shvetha Sankaran for proofing the manuscript.

