## Supplementary Figures for "*ctQC* improves biological inferences from single cell and spatial transcriptomics data"

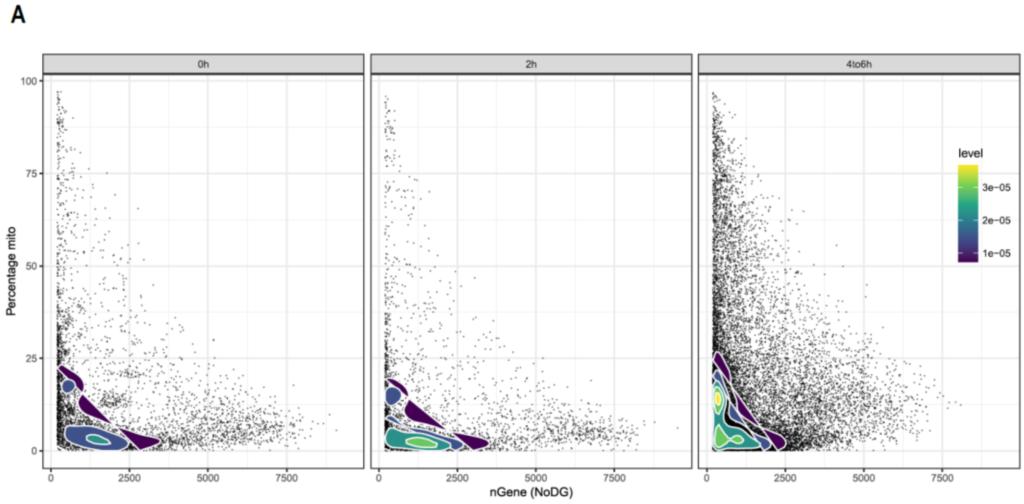


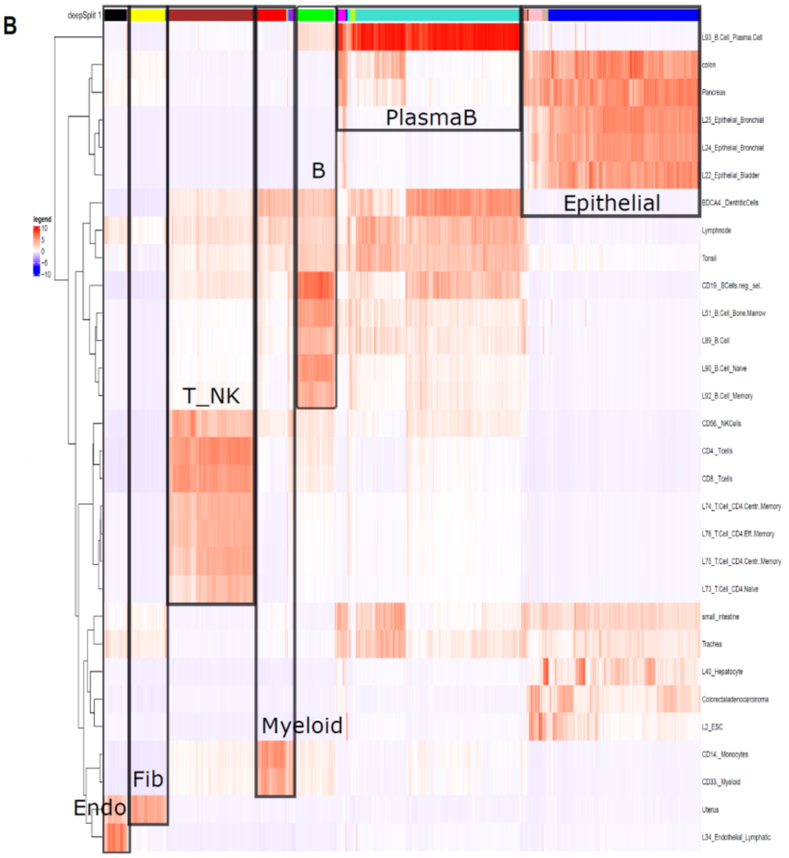


| | |
|---|---|
| Epithelial | 7684 |
| Fibroblast | 721 |
| Endothelial | 698 |
| PlasmaB | 7819 |
| B cell | 1003 |
| Myeloid | 1624 |
| T & NK | 4040 |

**Supplementary Figure S1:** (**A**) QC metrics scatter plot for all the cells (N= 31,619 cells) from the CRC timepoint dataset. Cells are grouped by timepoints. (**B**) Heatmap from RCAv2 shows annotation of clusters based on its global reference panel annotation. Table shows number of cells per annotated cell types.

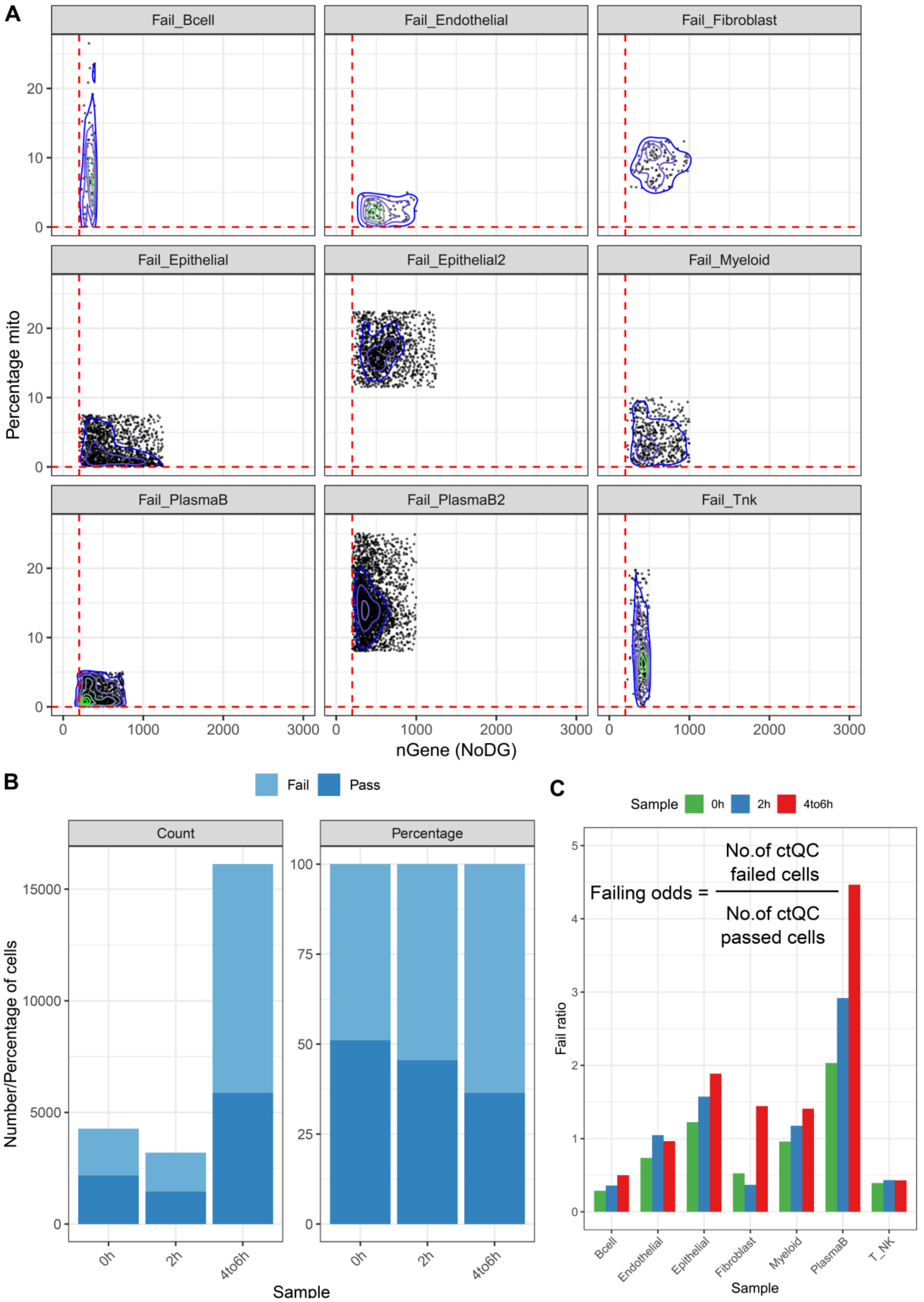


**Supplementary Figure S2:** (**A**) Cells that failed the cell type-specific QC cutoffs are plotted in QC metrics scatter plot. (**B**) Bar plot shows proportion of cells that passed or failed *ctQC* across different timepoints. (**C**) Failing odds ratio plotted for broad cell types across different profiled timepoints.

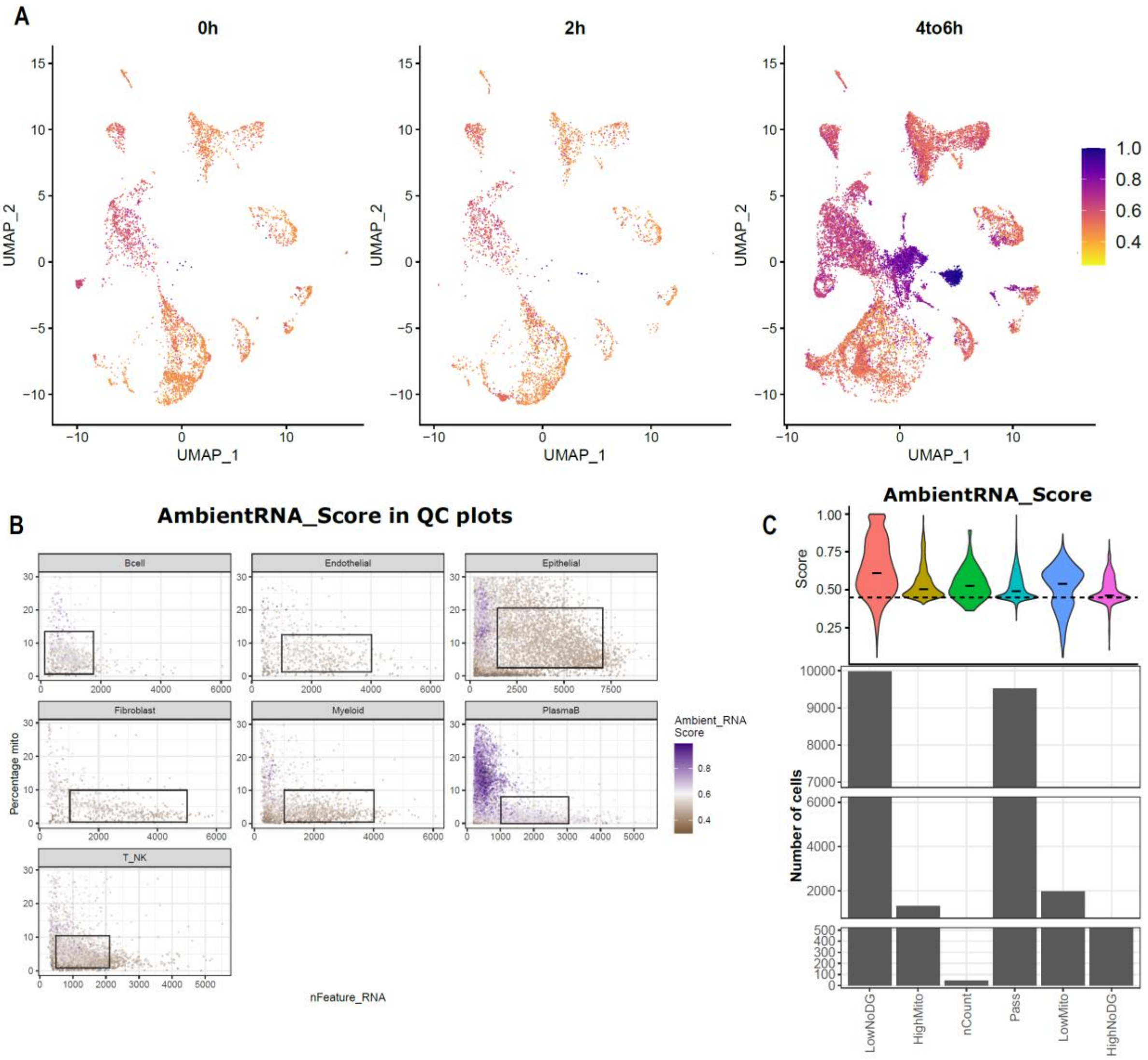


**Supplementary Figure S3:** (**A**) UMAP coloured by ambient RNA score across three timepoint samples. (**B**) Cell type wise QC metrics scatter plot where they are coloured by ambient RNA levels (module score). (**C**) The cells rejected by *ctQC* are classified into distinct groups according to the QC metric values they did not meet. Bar plot shows number of cells in each of these category and violin plot shows distribution of ambient RNA score for these categories.

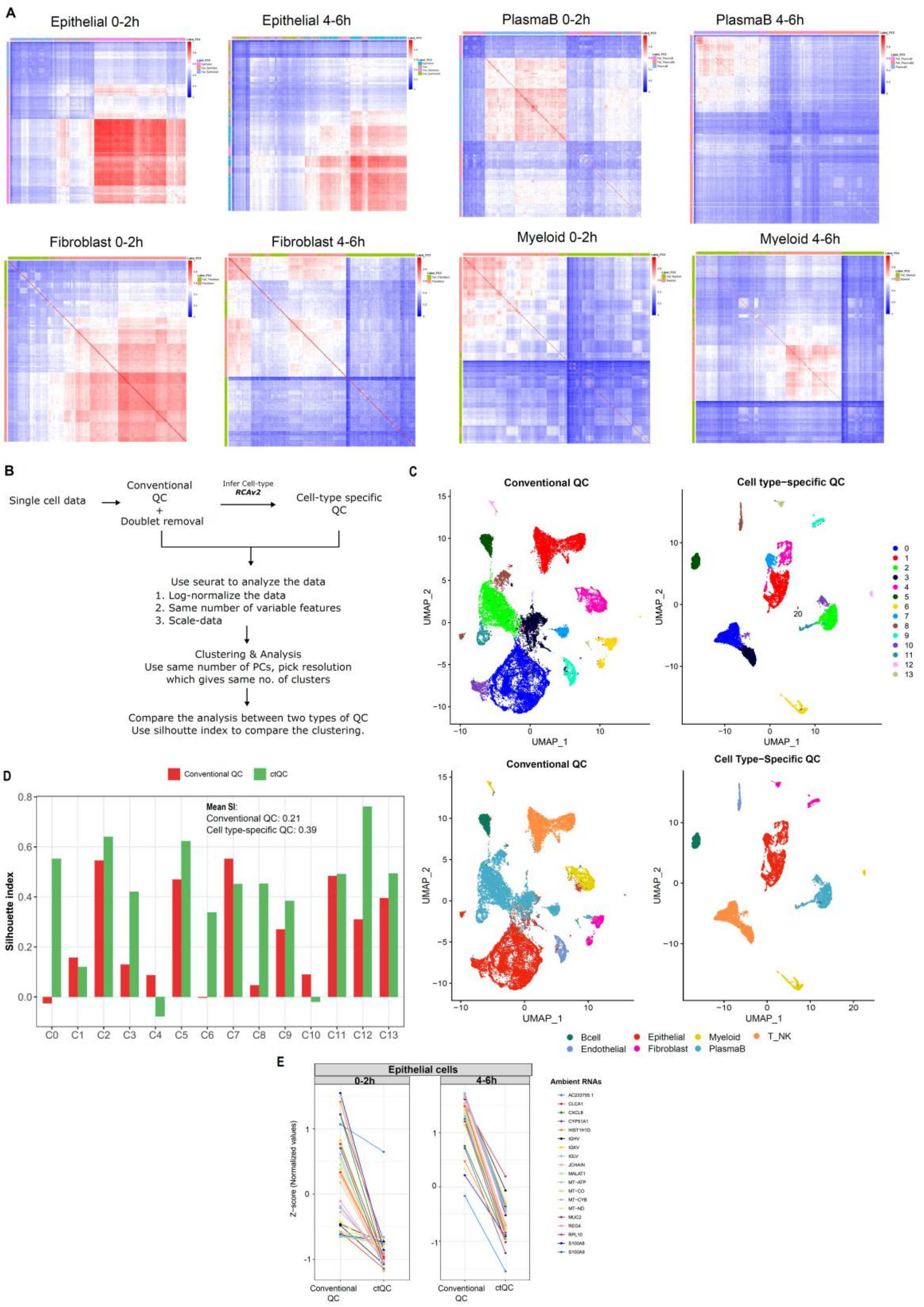


**Supplementary Figure S4:** (**A**) Cell to cell correlation (transcriptional similarity) is plotted for cells that passed and failed *ctQC* for individual broad cell lineage. Cells from 0-2h and 4-6h samples were pooled for this analysis. (**B**) Downstream single cell analysis pipeline used in this study to compare cells post *ctQC* and conventional QC. (**C**) Seurat annotated clusters and broad cell type label is plotted in UMAP embedding space post conventional (N= 23589) and cell type-specific QC (N= 9518). (**D**) Bar plot shows silhouette index calculated per cluster post conventional and *ctQC*. (**E**) Line plot shows scaled expression of top 20 ambient RNA genes (union of SoupX & cellbender predicted genes) in epithelial cells across 0-2h and 4-6h samples.

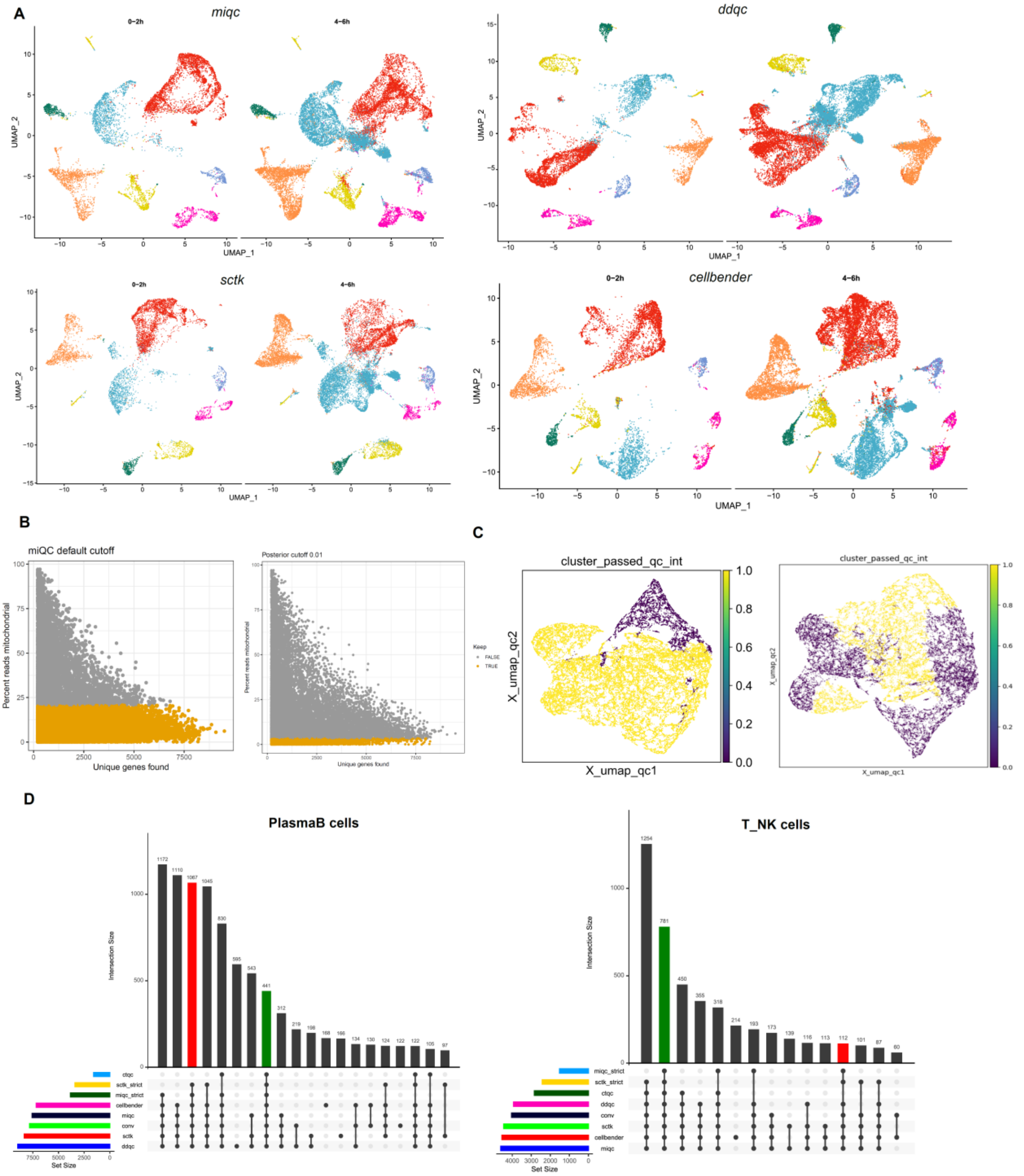


**Supplementary Figure 5:** (**A**) Cells retained by *miQC*, *sctk*, *ddqc* and *cellbender* with default QC cutoff were analysed similarly and UMAP are coloured by broad cell type label and split across timepoint. (**B**) Cells that passed default and stricter cutoff of *miQC* are coloured orange in QC metric plot. (**C**) Cells that passed default and stricter cutoff of *sctk* are highlighted in the umap where cells are clustered based on QC parameters. Left – default & Right – strict parameters. (**D**) Upset plot showing overlap of cells retained by different methods (at varying strictness). Green bar highlights cells retained by all the methods. Red bar highlights low-quality cells discarded by ctQC while retained by all the methods (varying strictness).

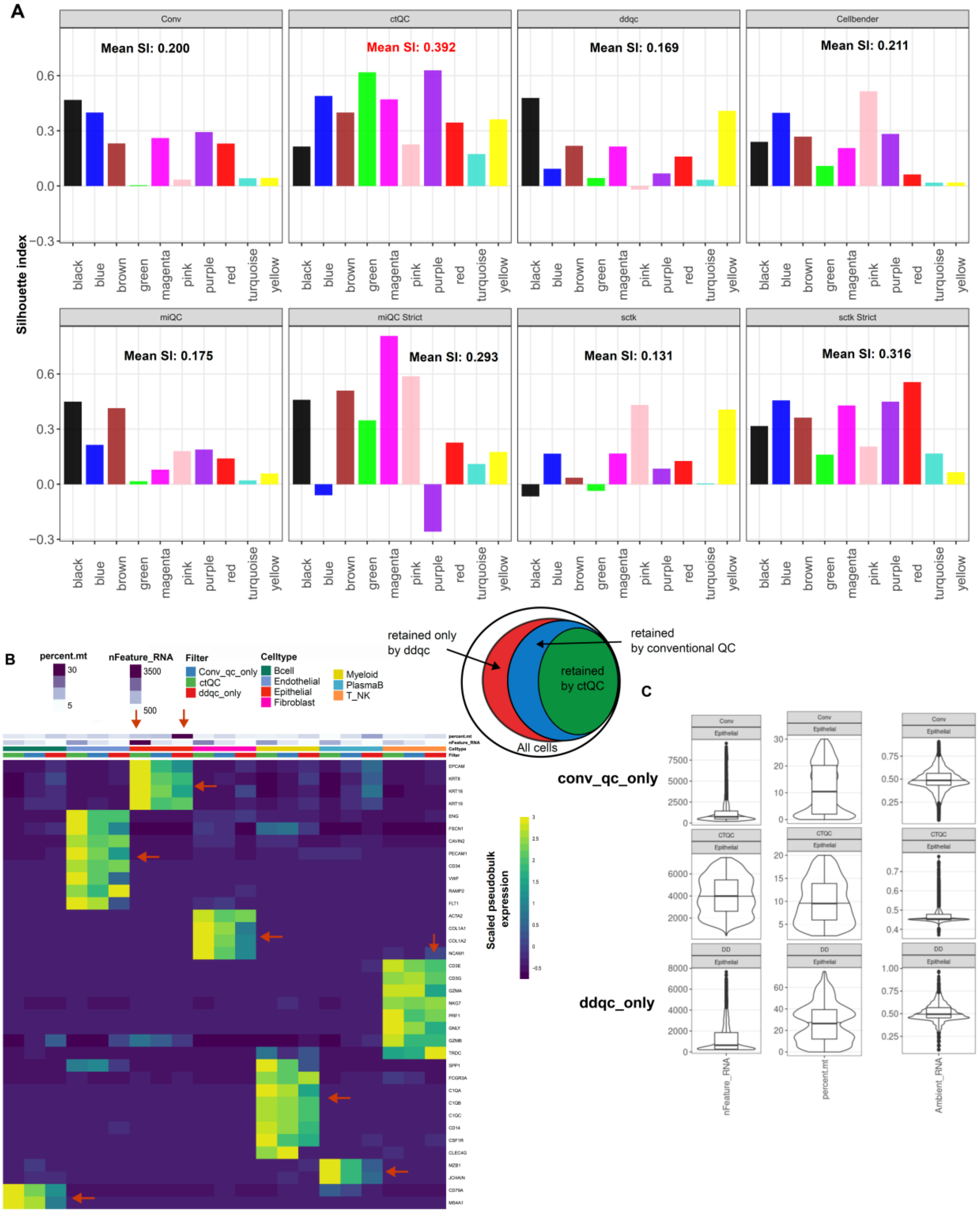


**Supplementary Figure S6:** (**A**) Silhouette index calculated for clusters derived from CRC timepoint data using different QC pipelines tested in this study. Clusters derived from RCAv2 is used here to calculate SI. (**B**) Heatmap shows expression levels of major cell type markers for cells that were retained exclusively by *ddqc* (N=4005), conventional(N=14071) and *ctQC* (N= 9518). Gene complexity (nFeature_RNA), percentage of reads mapping to mitochondrial genes, broad cell type annotations and QC groups are plotted as column annotation. Red arrows highlight cells exclusively retained by *ddqc*. (**C**) Distribution of nFeature_RNA, percent.mt and ambientRNA_score for epithelial cells across the three categories. Cells exclusively retained by *ddqc* has low NODG, high percent.mt and ambient RNA scores.

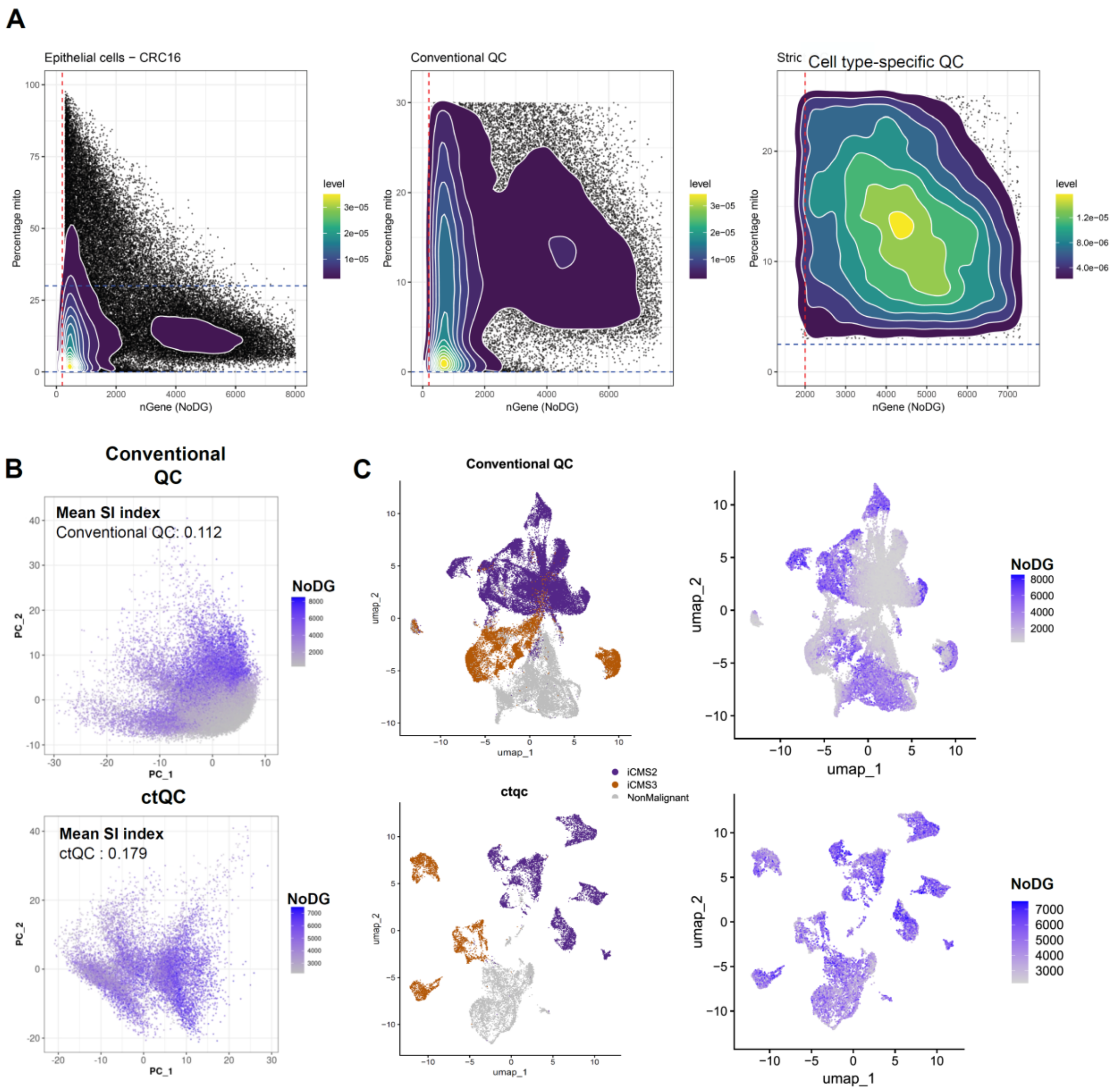


**Supplementary Figure S7:** (**A**) QC metric plot for all the epithelial cells from CRC-SG1 cohort, post conventional and *ctQC*. (**B**) PCA plot coloured by NODG for cells post conventional and cell type-specific QC. (**C**) UMAP coloured by CRC tumour intrinsic molecular stratification and by NODG.

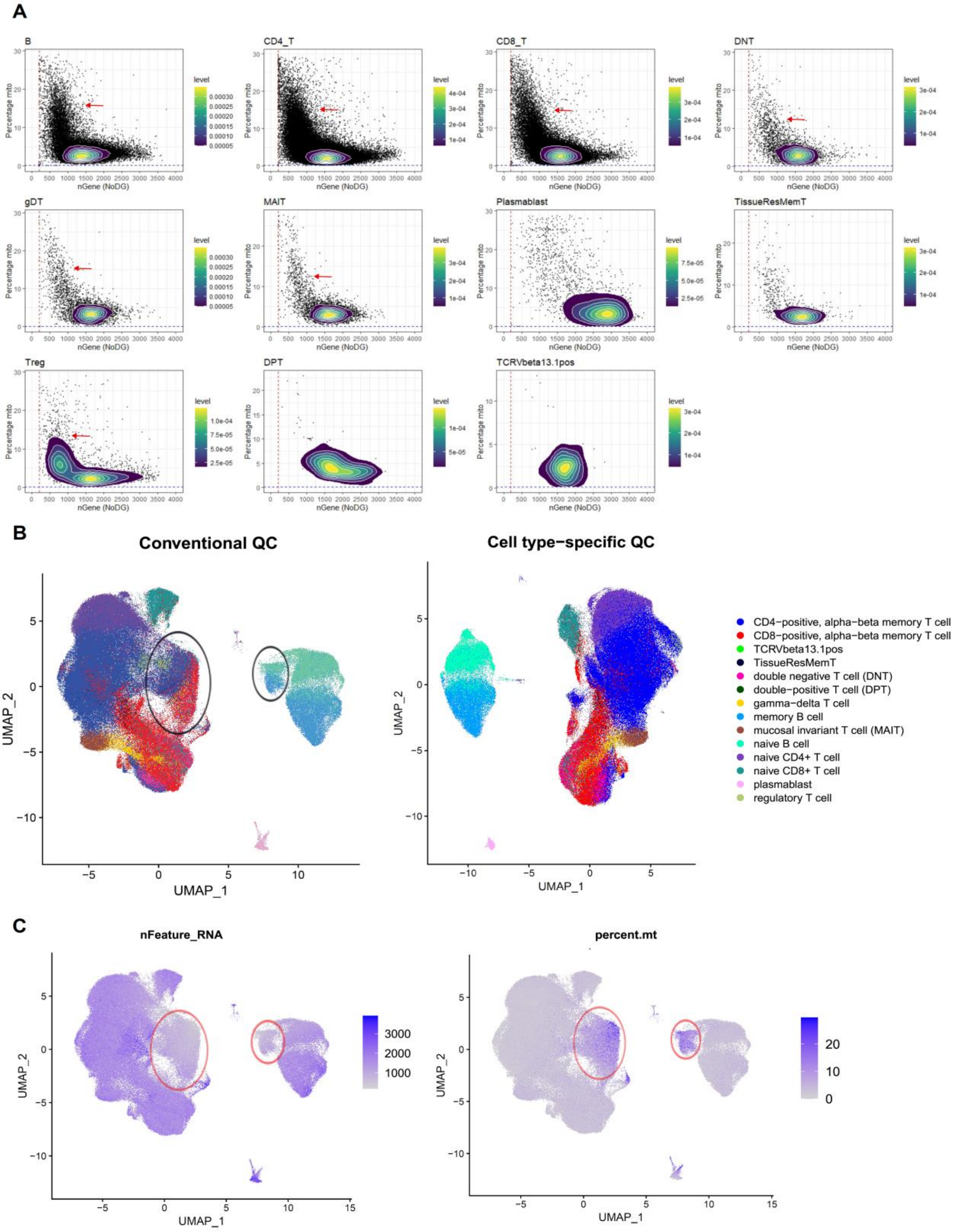


**Supplementary Figure S8:** (**A**) Cell type wise QC metric scatter for all adaptive immune cells post conventional QC. Cell type labels are adapted from authors annotations. Red arrow indicates the mode where most of the low-quality cells are enriched. (**B**) Adaptive immune cells post conventional and ctQC are shown in UMAP space and coloured by broad cell type labels and by QC parameters (NODG and percent.mt) in (**C**). Circled region has prominent signature of low-quality cells.

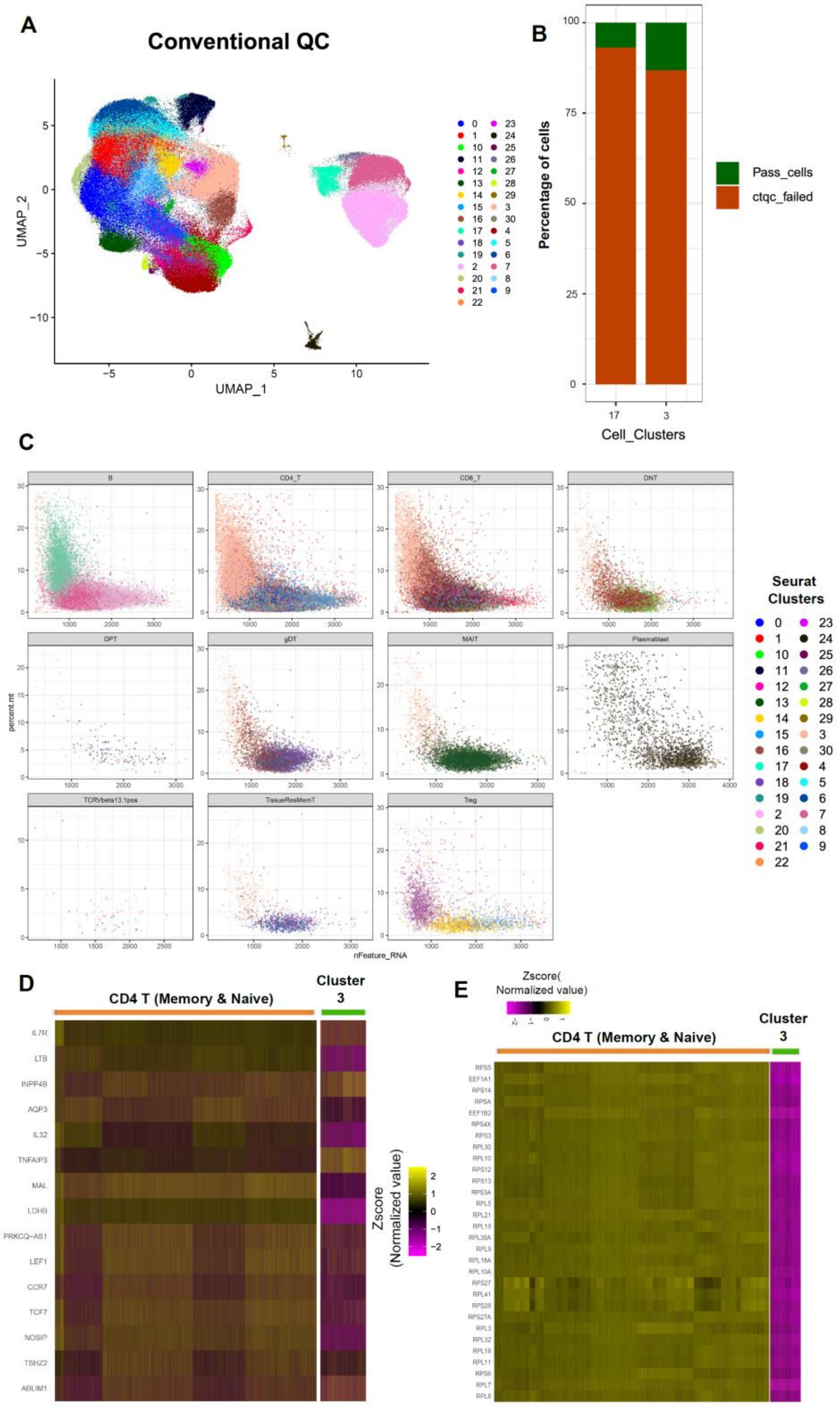


**Supplementary Figure S9:** (**A**) Adaptive immune cells post conventional QC are coloured by seurat clusters in UMAP space. (**B**) Stacked bar plot represents percentage of cells that failed *ctQC*. Cluster 17, 3 represent low-quality cluster from B and CD4 T. (**C**) QC metrics scatter plot of cells post conventional QC that are grouped by author cell type annotations and cells are coloured by Seurat clusters. (**D**) Heatmap showing expression of CD4_T cell type markers for cells that were retained & discarded by *ctQC*. (**E**) Heatmap showing expression of top 30 down-regulated genes in cluster 3 relative to other CD4 T cells.

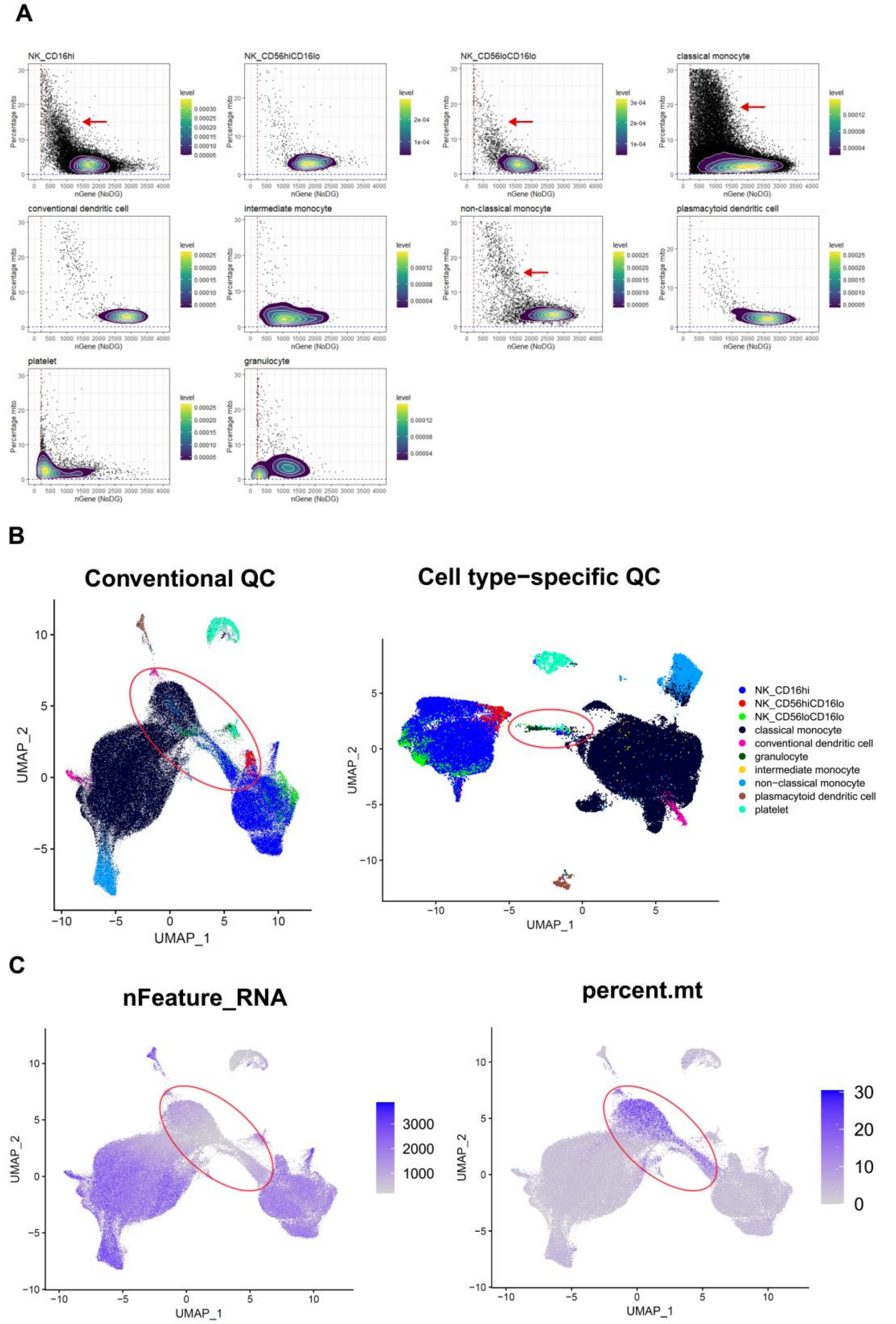


**Supplementary Figure S10:** (**A**) Cell type wise QC metric scatter for all innate immune cells post conventional QC. Cell type labels are adapted from original study. Red arrow indicates the mode where most of the low-quality cells are enriched. (**B**) Innate immune cells post conventional and *ctQC* are shown in UMAP space and coloured by broad cell type labels and by QC parameters (NODG and percent.mt) in (**C**). Circled region has prominent signature of low-quality cells and bridging cells between CD16hiNK and classical monocytes.

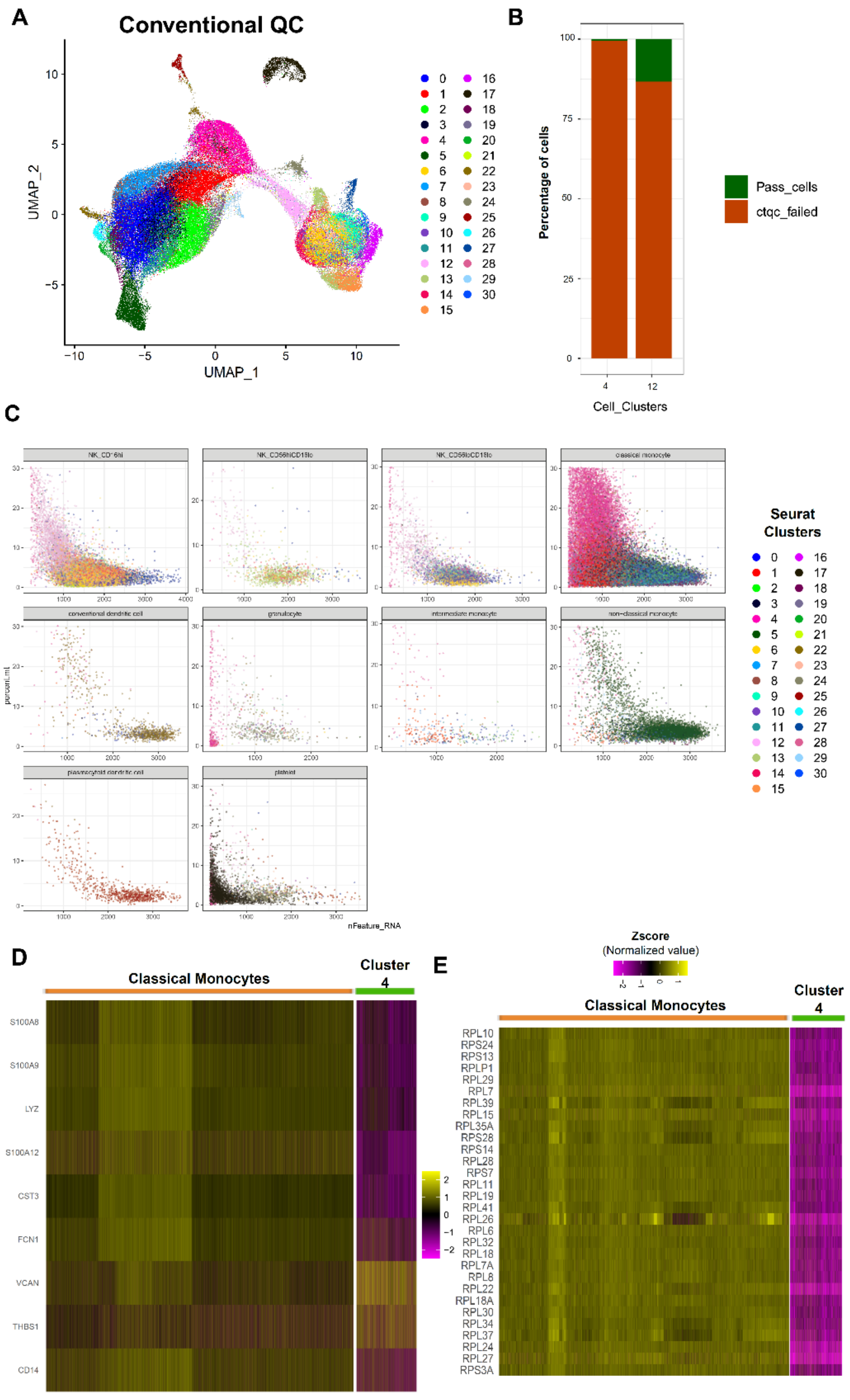


**Supplementary Figure S11:** (**A**) Innate immune cells post conventional QC are coloured by seurat clusters in UMAP space. (**B**) Stacked bar plot represents percentage of cells that failed *ctQC* cutoffs. Seurat cluster 4, 12 represent low-quality cluster corresponding to classical monocyte & CD16hiNK. (**C**) QC metrics scatter plot of cells post conventional QC that are grouped by author cell type annotations and cells are coloured by Seurat clusters. (**D**) Heatmap showing expression of classical monocyte markers for cells that were retained & discarded by *ctQC*. (**E**) Heatmap showing expression of down-regulated genes in cluster 4 relative to other monocyte cells.

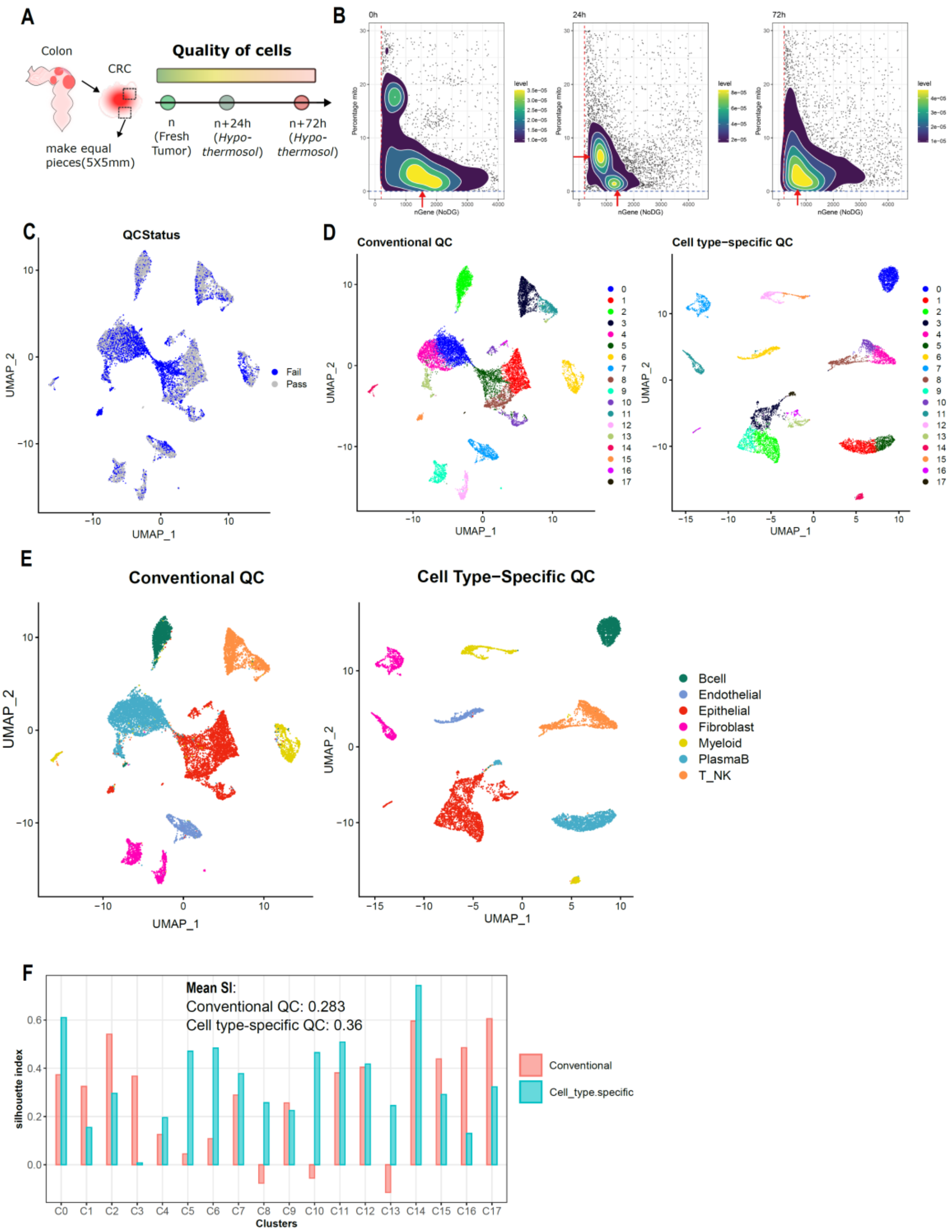


**Supplementary Figure S12:** (**A**) Schematic depicts experimental design where samples from CRC patients were collected and stored in hypothermosol for 24h & 72h and subjected to scRNA seq. (**B**) QC metrics scatter plot of all cells post conventional QC across different timepoints. Red arrow indicates the mode of the distribution. (**C**) UMAP highlights cells failed *ctQC* cutoffs and were retained by conventional QC cutoffs. (**D**) Post conventional and cell type-specific QC, cells were analysed using Seurat and final UMAP embedding of annotated clusters are shown and broad cellular annotations are shown in (**E**). (**F**) Bar plot shows silhouette index calculated per cluster wise post conventional and *ctQC*.

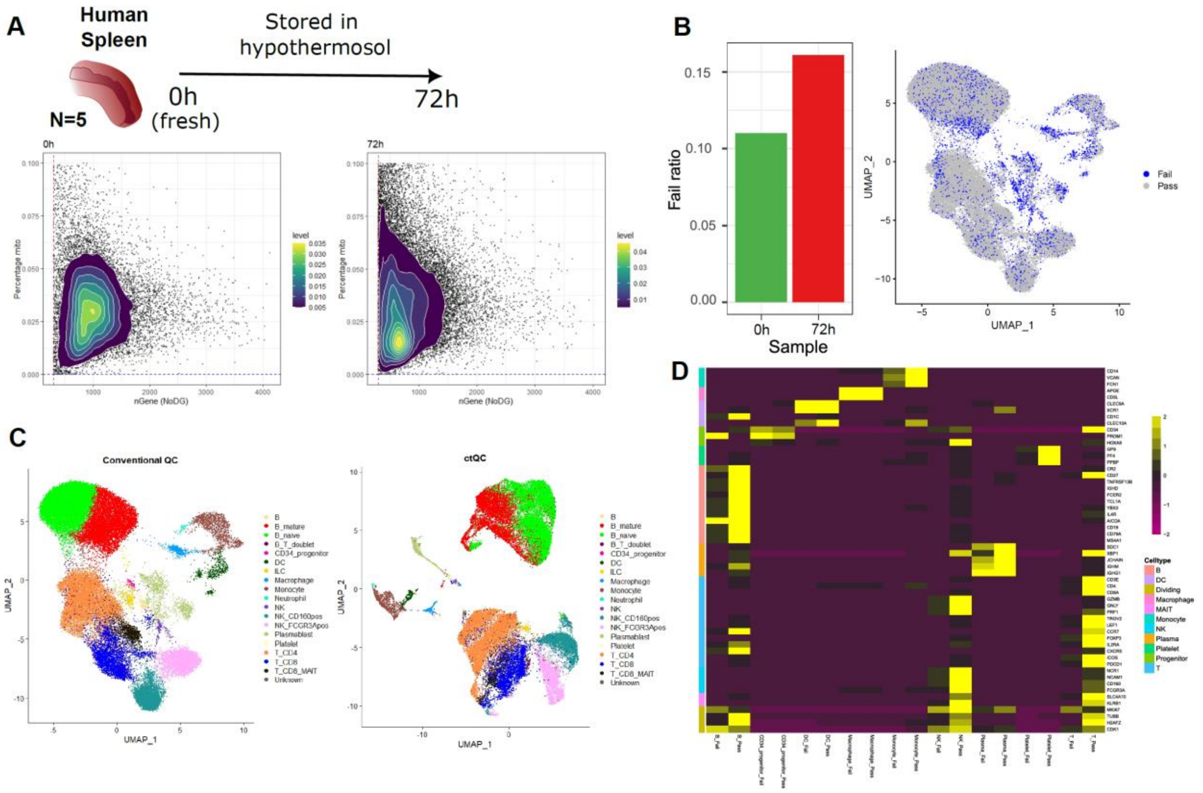


**Supplementary Figure S13:** (**A**) Human spleen scRNA-seq data is collected from fresh (0h) and 72h of storage in preservative medium (hypothermosol). All the sequenced cells are shown in QC metric scatter plot. (**B**) Fail ratio plotted for the two profiled time points. *ctQC* failed cells are highlighted in blue colour in UMAP space. (**C**) Comparing the cell type separation in UMAP space post conventional and cell type-specific QC. (**D**) Heatmap shows expression levels of major cell type markers for cells that passed and failed *ctQC*.

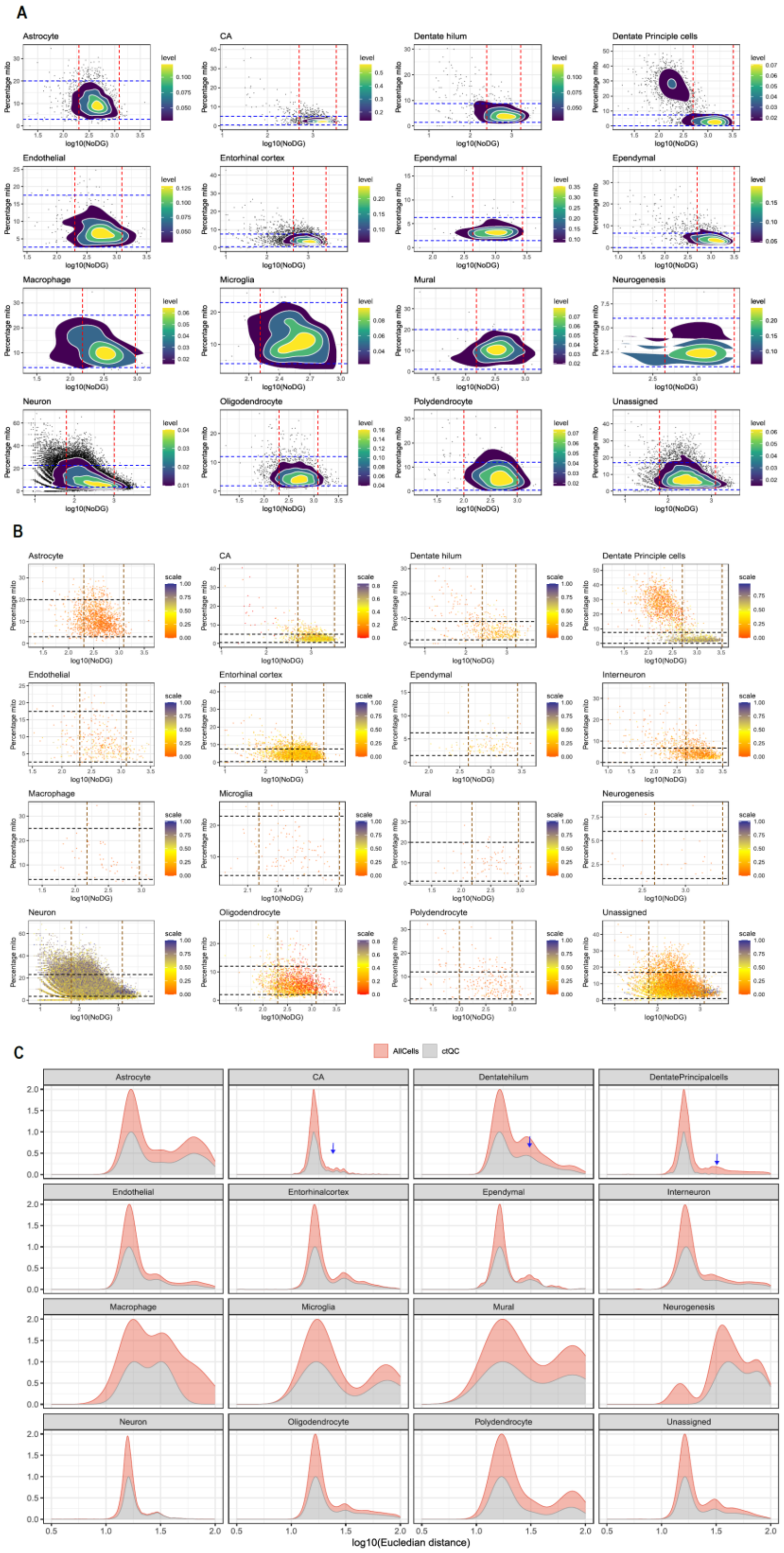


**Supplementary Figure S14:** (**A**) Cells from hippocampus slide-seq data were grouped into major cell types based on authors annotation and cells are visualized in QC metric space. Red & blue lines indicate the cut-offs used for *ctQC*. (**B**) Neighbourhood score calculated for hippocampus slide-seq data is overlayed in QC metrics scatter plot. The dotted lines indicate the cut-off used for *ctQC*. (**C**) Closeness score calculated for pucks post conventional & *ctQC* are plotted as density plot.

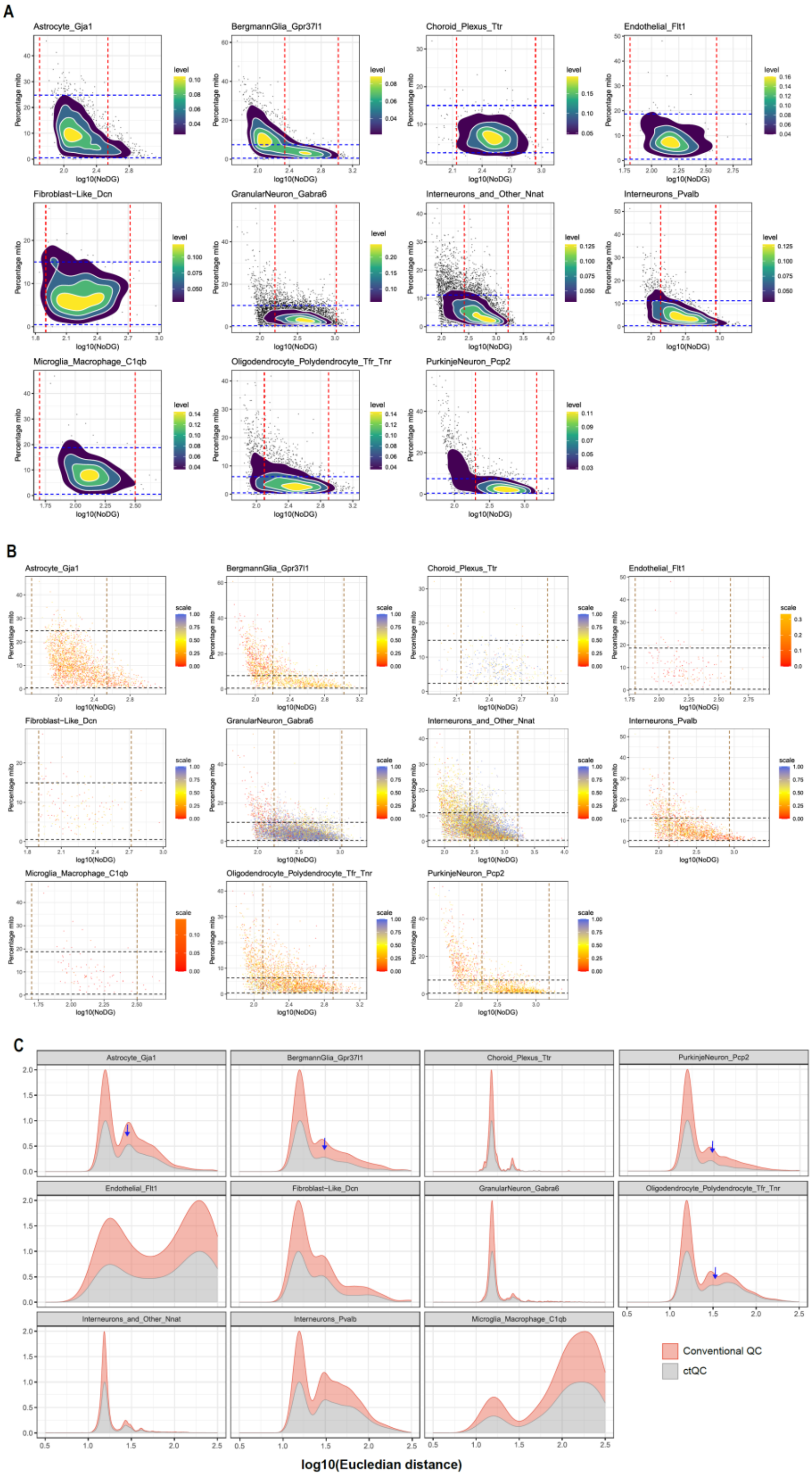


**Supplementary Figure S15:** (**A**) Cells from hippocampus slide-seq data were grouped into major cell types based on authors annotation and cells are visualized in QC metric space. Red & blue lines indicate the cut-offs used for *ctQC*. (**B**) Neighbourhood score calculated for all the cells in cerebellum slide-seq data is overlayed in QC metrics scatter plot. The dotted lines indicate the cut-off used for *ctQC*. (**C**) Closeness score for pucks post conventional & *ctQC* are plotted as density plot.

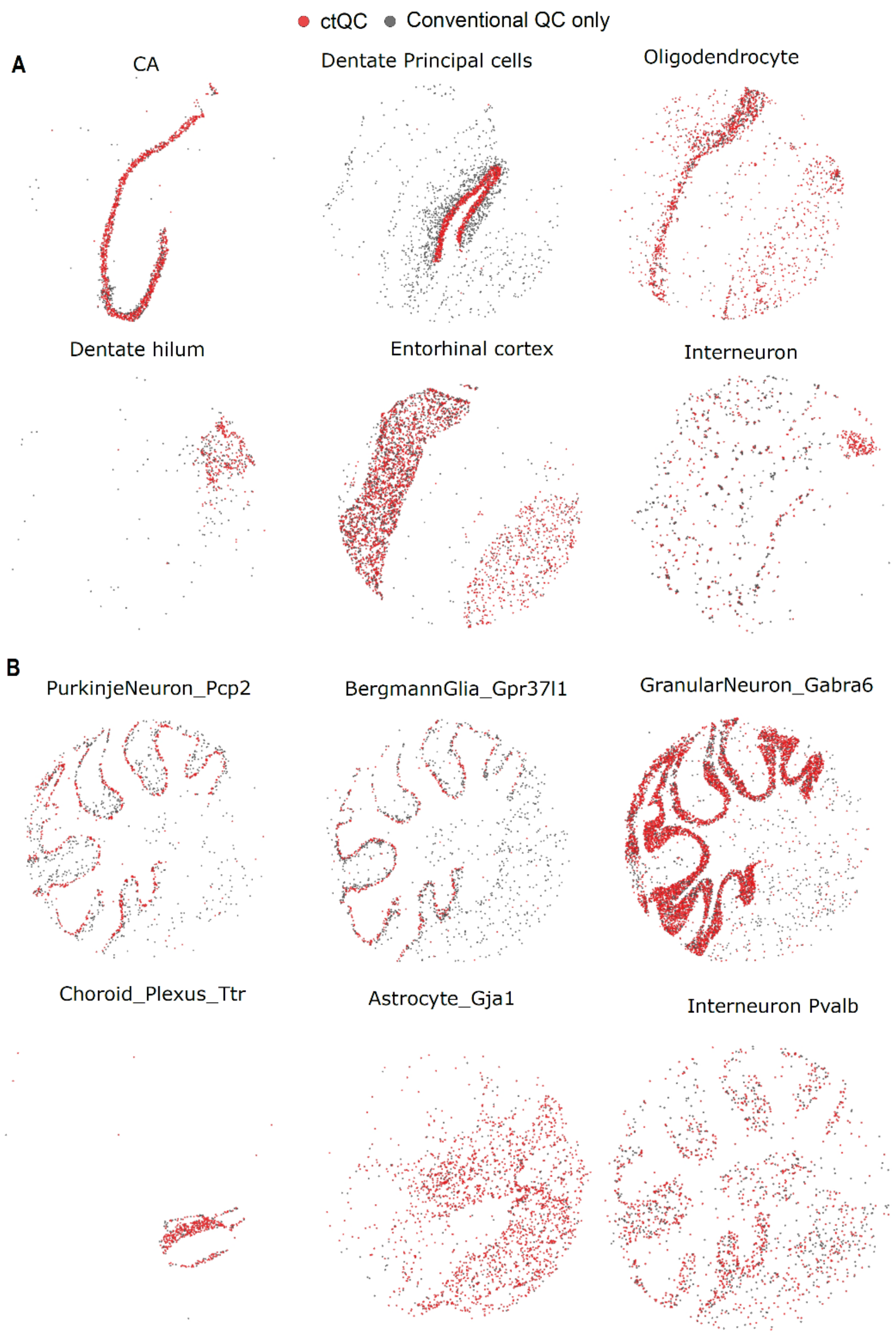


**Supplementary Figure S16:** (**A**) Spatial distribution of pucks retained post *ctQC* highlighted as red for various cellular subtypes (based on authors annotations) for hippocampus dataset. Pucks coloured in grey are cells exclusively retained by conventional QC. (**B**) is similar to (**A**) where pucks retained by *ctQC* are highlighted in red colour for cerebellum dataset.
